# APNet, an explainable sparse deep learning model to discover differentially active drivers of severe COVID-19

**DOI:** 10.1101/2024.01.11.575161

**Authors:** George I. Gavriilidis, Vasileios Vasileiou, Stella Dimitsaki, George Karakatsoulis, Antonis Giannakakis, Georgios A. Pavlopoulos, Fotis Psomopoulos

## Abstract

**Motivation:** Computational analyses of plasma proteomics provide translational insights into complex diseases such as COVID-19 by revealing molecules, cellular phenotypes, and signaling patterns that contribute to unfavorable clinical outcomes. Current *in silico* approaches dovetail differential expression, biostatistics, and machine learning, but often overlook nonlinear proteomic dynamics, like post-translational modifications, and provide limited biological interpretability beyond feature ranking.

**Results:** We introduce APNet, a novel computational pipeline that combines differential activity analysis based on SJARACNe co-expression networks with PASNet, a biologically-informed sparse deep learning model to perform explainable predictions for COVID-19 severity. The APNet driver-pathway network ingests co-expression and classification weights to aid result interpretation and hypothesis generation. APNet outperforms alternative models in patient classification across three COVID-19 proteomic datasets, identifying predictive drivers and pathways, including some confirmed in single-cell omics and highlighting under-explored biomarker circuitries in COVID-19.

**Availability and Implementation:** APNet’s R, Python scripts and Cytoscape methodologies are available at https://github.com/BiodataAnalysisGroup/APNet

**Supplementary information:** Supplementary information can be accessed in Zenodo (10.5281/zenodo.10438830).

## 1. Introduction

Human plasma is a vital clinical specimen encompassing a broad spectrum of proteins, including tissue markers, immunoglobulins, transcription factors, kinases, metabolites, and secreted factors (Eldjarn et al., 2023; Zhong et al., 2021). With the advent of high-throughput technologies (-omics), the human plasma proteome has become a focal point for discovering novel biomarkers and therapeutic targets for complex diseases. This has been especially the cases with severe COVID-19, a condition besetting many patients infected with the SARS-CoV-2 coronavirus (Babačić et al., 2023). Plasma proteomics have provided significant biological insights into the immunopathology of severe COVID-19, which is characterized by the inflammatory “cytokine storm”, Acute Respiratory Distress Syndrome (ARDS), PANoptosis-induced cell death, and multiorgan failure (Diamond and Kanneganti, 2022). Plasma proteomics has also been explored in long-COVID-19 syndromes and vaccine response variations (Liang et al., 2023).

Many studies have measured plasma proteomics using Olink Proximity Extension Assay (PEA) in COVID-19 research due to this technology’s specificity, scalability and multiplexing benefits (Wik et al., 2021). In our recent work, we assessed pertinent Machine Learning models applied in these high-dimensional datasets like Random Forest, Gradient Boosted Decision Tree, XGBoost, Extra Tree classifiers, Logistic regression, Lasso Logistic regression, Support Vector Machine (SVM), and Deep Learning (DL) (e.g., AutoGluon-Tabular). Some models exhibited eXplainable AI (XAI) features by deploying Shapley additive explanation (SHAP) values, the minimal-optimal variables method or a random forest explainer. In the same work, we managed to dovetail an explainable, computational pipeline to benchmark a wide assortment of ML tools on predicting COVID-19 severity from Olink plasma proteomics which revealed Multi-Layer Perceptron (MLP) as the highest-performing algorithm (Dimitsaki et al., 2023).

However, most of the above studies can partially approximate proteomic non-linear dynamics (e.g., post-translational modifications, protein co-expression networks, complex formation, and subcellular localization), thus missing signaling proteins that may drive critical COVID-19 pathways. Moreover, these studies’ ML/DL findings often lack extensive external validation in large independent datasets, while their biological explainability is usually restricted to mere feature ranking (Paul et al., 2023) (Dimitsaki et al., 2023).

Acknowledging these challenges, we introduce Activity PASNet (APNet) in this manuscript. This computational DL pipeline initially uses the SJARACNe data-driven network algorithms to uncover disease drivers prioritized based on “activity,” an aggregate metric of their capacity to regulate their transcriptional targets non-linearly (Ding et al., n.d.; Dong et al., 2023). These drivers can be overt (differentially expressed and possibly active) or “hidden” (differentially active but not expressed). Next, APNet feeds these drivers into Pathway-Associated Sparse Deep Neural Network (PASNet) (Hao et al., 2018), which incorporates biological priors as hidden layers to ultimately deliver interpretable clinical classifications, validated by the eXplainable AI component of SHAP values. Finally, APNet facilitates the analysis of SJARACNe co-expression networks, equipped with the weights from the DL classification task, to streamline data exploration and the formation of mechanistic hypotheses for further biological investigation.

We extensively trained, tested, and validated APNet on activity matrices from 3 distinct Olink plasma proteomic datasets (MGH, Mayo, Stanford) (Byeon et al., 2022; Feyaerts et al., 2022; Filbin et al., 2021). APNet managed to pinpoint ground-truth drivers of severity, predicted new proteomic markers with potential theranostic potential (some of which were traced to circulating PBMCs through scRNA-seq analysis), outperformed alternative ML/DL models in demarcating severe COVID-19 cases and enabled the inference of a potential signaling network from predictive factors in the liver of individuals with severe COVID-19.

## 2. Materials and methods

### 2.1 APNet overview

APNet is a modular pipeline (Figure 1) which aims to facilitate the discovery of novel predictive drivers of severe clinical outcomes and to facilitate the formulation of mechanistic hypotheses. In this present work, we considered cases experiencing severe and non-severe COVID-19.

**Figure 1.**
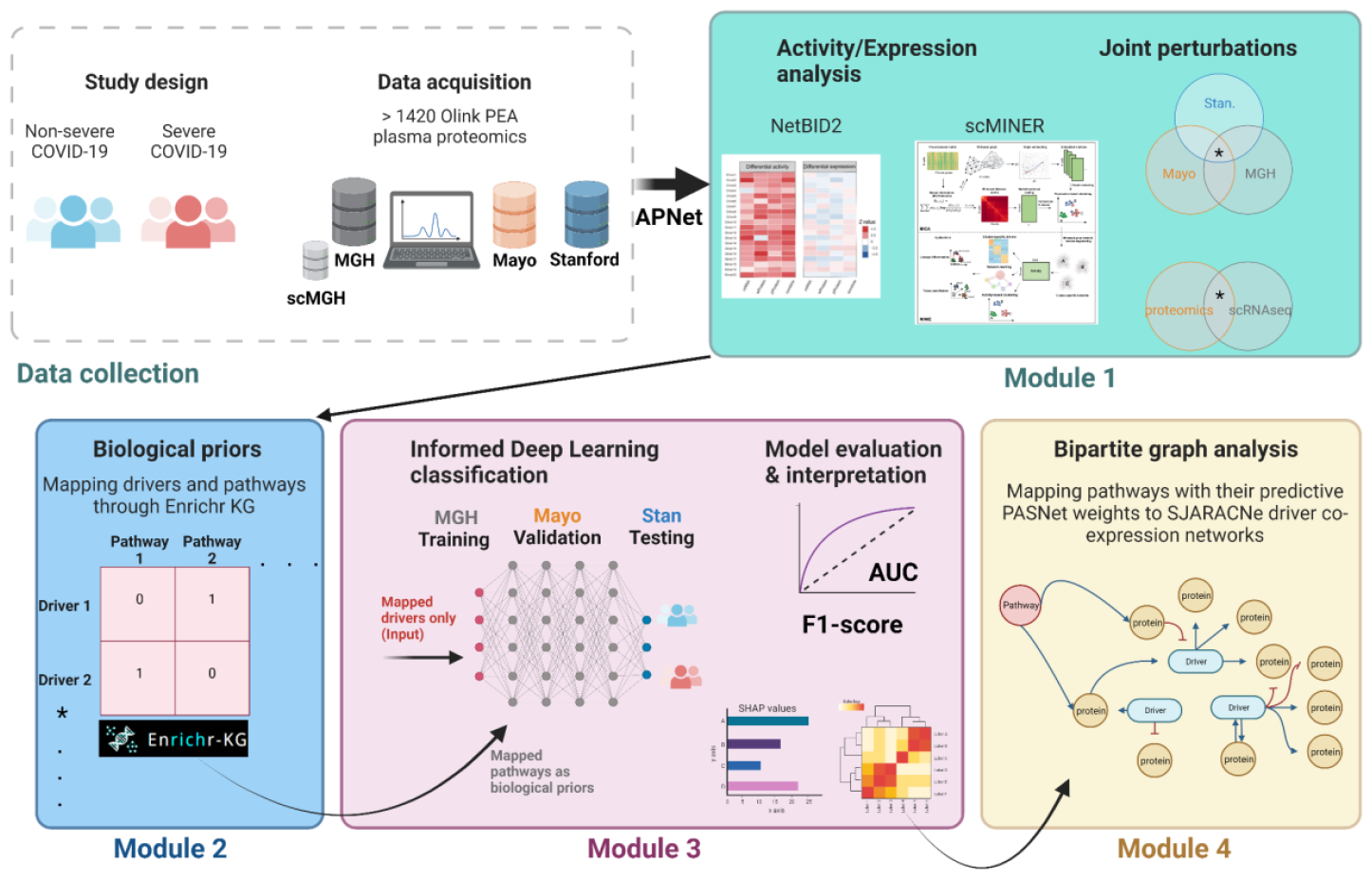
APNet workflow, as implemented in the herein COVID-19 multi-omic study to discover predictive drivers of severity. Image made using the Biorender toolkit.

### 2.2 Brief description of APNet modular architecture

#### 2.2.1 Module 1- Differential activity analysis for drivers of COVID-19 severity

In this module, conversions of expression values to activity values for plasma proteomics were accomplished with NetBID2 (Dong et al., 2023) toolkit whereas for scRNA-seq data with scMINER toolkit (Ding et al., 2023). For the plasma proteomics, we applied the NetBID2 algorithm, which reverse-engineers context-specific interactomes and integrates network activity inferred from large-scale multi-omics data, empowering the identification of hidden drivers that traditional analyses cannot detect. By leveraging the MSigDB database, we compiled distinct lists of Transcription Factors (TF) and signaling molecule proteins. Separate TF and signaling molecule networks were constructed using SJARACNe. These networks featured drivers (hubs) connected to their targets through protein-protein interactions, derived from their expression patterns.

To calculate the activities of driver proteins in each dataset based on protein expression, we employed the “cal.Activity” function in NetBID2. The weighted mean activity of a driver candidate protein (Driver{i}) in sample s, was computed using the following equation:

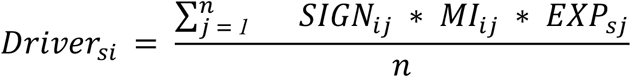

Here, the NPX count proteomics matrix, EXP{sj} represented the expression value of gene j in sample s, MI{ij} indicated the mutual information between master regulator protein i and its target protein j, and SIGN{ij} was the sign of the Spearman correlation between protein i and its target protein j. The total number of targets for DRIVER i was denoted by n.

Differential activities were then computed for Severe and Non-Severe Status across the three datasets, by using the “getDE.BID.2G” function, allowing us to identify genes exhibiting distinct regulatory patterns in response to severity variations, through Bayesian model.

Also, we deployed the scMINER workflow, based also on SJARACNe, to discover severity drivers in MGH scRNAseq data. For both differential expression and differential activity, the function get.DA was performed by using the SCT matrix and activity matrix, respectively. Data visualisation for single-cell analysis was performed through the Seurat pipeline (4.3.0).

#### 2.2.2 Module 2- Driver-pathway mapping

To prepare input data for the biologically explainable PASNet DL model on Module 3, joint differentially active drivers of severity from the three Olink studies were mapped to biological pathways using the Enrichr KG (Evangelista et al., 2023). 30 pathways from each of the following resources were leveraged (KEGG, Reactome, GO:BP and Wikipathways 2021) for the commonly decreased and increased drivers of severity separately. Drivers were mapped to the retrieved pathways in a binary fashion with 0s and 1s, i.e. when a driver was participating in the gene set of a pathway it was assigned the value of 1 and vice versa.

#### 2.2.3 Module 3-Deep Learning classification of Severe COVID-19 cases with biological explainability

The findings from Modules 1 and 2 served as input for Module 3, where a sparse neural network model called PASNet was used to predict COVID-19 severity. The model was trained on MGH data, validated on Mayo and tested on Mayo and Stanford datasets. A separate model was trained and tested using scMGH data. Model performance was evaluated using Area Under the Curve (AUC) and F1-scores, along with ROC curve analysis. The PASNet training phase is expressed through the following equations.

Sparsity of the PASNet sub-network function: *h*^(*l*+*1*)^ = *a*((*W*^(*l*)^ * *M*^(*l*)^)*h*^(*l*)^ + *b*^(*l*)^)

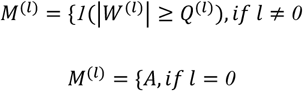

where

*Q*^(*l*)^ is the S-th percentile of |*W*^(*l*)^| if *l* ≠ *0*

M: mask matrix for each layer

*l*:layer

W: weight matrix

b: vias vector

Cost-sensitive learning for imbalanced data: 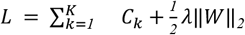

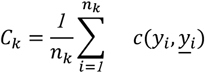

Thus, the weights and biases on the *l*-th layer are updated by:

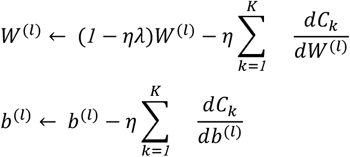

where

*C*_*k*_:mean error on the class k

*y*_*i*_:ground truth

<u>*y*_*i*_</u>:prediction

*n*_*k*_:number of samples in the class k

*L*: total cost

*c*(.):cost function (e.g., cross-entropy loss)

*λ*: regularization hyperparameter

*η*:learning rate

Biological explainability of the whole sparse DL model is predicated in the combination of Shapley values (SHAP) and the driver-pathway mappings that PASNet architecture offers, assigning learning weights.

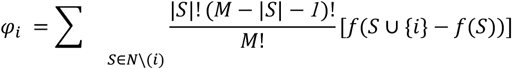

*N*: is the set of all input features

*i*: feature

*f*: model

*M*: is the number of features

#### 2.2.4 Module 4- Bipartite graph analysis

This final APNet module leverages the SJARACNe co-expression networks from each study for the joint differential active drivers and augments it by connecting drivers to pathways based on Module 2. The weights of driver-driver edges contain the Mutual Information (MI) metric and the Spearman correlation coefficient (positive values signify activation, negative values the opposite), amongst other metrics. Driver-pathway edges contain as weights the PASNet-weights that PASNet learned during training-testing tasks from Module 3. Our study used Cytoscape to perform network visualization, basic analysis for network statistics and centrality metrics (Betweenness Centrality algorithm), dimensionality reduction using tSNE (cluster signal propagation simulation (OCSANA+) and analysis for shortest paths (PathLinker tool)(Gil et al., 2017; Marazzi et al., 2020).

OCSANA+ is a Cytoscape application that analyses the structure of large-scale complex networks. It identifies nodes that drive the system towards a desired long-term behavior and ranks the combinations of interventions that are likely to be more effective. Additionally, it estimates the effects of perturbations in signaling networks. We used the Signal Flow Analysis (SFA) feature of OCSANA+ to simulate signal propagation. The SFA algorithm estimates the signal flow in a signaling network by analyzing the topological information. It employs a linear difference equation that considers a node’s previous activity, the effect and influence of incoming edges, and the initial activities of the node. The algorithm focuses on the information conveyed by a series of biological interactions represented in a signaling network (Marazzi et al., 2020).

PathLinker is a Cytoscape app based on an algorithm reconstructing interactions in a signaling pathway. It requires a directed network, a set of sources, and a set of targets as inputs. The algorithm computes the k best-scoring loopless paths and outputs the sub-network of the k best paths. The algorithm offers three choices for managing edge weights: (i) No weights: Path score is based solely on the number of edges in the path, and PathLinker identifies the k paths with the lowest scores, (ii) Additive edge weights: Path score results from the summation of edge weights, and PathLinker finds the k paths with the lowest scores in this scenario as well, (iii) Probabilistic edge weights: Common in protein interaction networks, where weight represents experimental reliability. PathLinker treats these weights as multiplicative, seeking the k paths with the highest cost, where the product of edge weights determines cost. Internally, PathLinker transforms each weight by taking the absolute value of its logarithm to map the problem to the additive case (Gil et al., 2017) (Figure 2).

**Figure 2.**
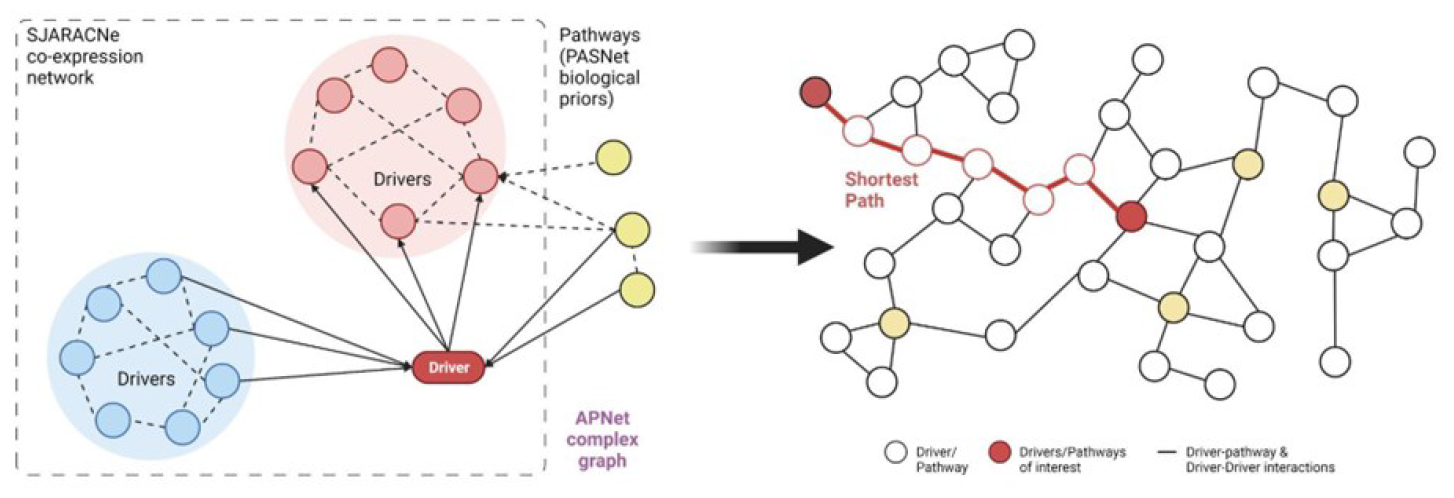
Outline of APNet complex graph and the ensuing analysis with shortest path algorithms for uncovering non-intuitive connections among drivers and pathways.

### 2.3 Technical benchmarking and bioinformatic validation based on COVID-19 prior knowledge

To benchmark APNet’s performance on patient classification from Olink plasma proteomics, we deployed the PASNet approach on original NPX values of Olink plasma proteomics and a Random Forest model on the transformed activity values. Firstly, for PASNet we used the expression values for training, validation and test, by using the count matrices of MGH, Mayo and Stanford, respectively. The count matrices were filtered by keeping only the common significant proteins across 3 datasets from Differential Expression analysis to perform the PASNet approach on expression data. Similarly to the APNet approach, pathways collected from EnrichR-KG, by using (KEGG, Reactome, GO:BP and Wikipathways 2021) for the commonly decreased and increased drivers of severity separately. PASNet used the count matrices across 3 datasets for training, validation, and test, by using MGH, Mayo, and Stanford respectively. Then for Random Forest, we used activity matrices from 3 datasets, by applying training, validation and test into MGH, Mayo and Staford, respectively.

Bioinformatic validation of the top 20 most predictive drivers for each experiment was pursued by mapping these drivers to the 9 curated networks regarding COVID-19 immunopathological hallmarks by SIGNOR (https://signor.uniroma2.it/covid/). Level 4 networks were obtained for each COVID-19 hallmarks and downstream processing was conducted in Cytoscape.

Finally, selective data mining for key drivers of interest was performed in the web tool https://www.covid19dataportal.org/.

### 2.4 DOME recommendations

The assembly of APNet was performed considering the recently published DOME recommendations, a set of community-wide recommendations for reporting supervised machine learning–based analyses applied to biological studies (see supplementary information) (Walsh et al., 2021) (Supplementary Material 1).

## 3. Results

### 3.1 Harmonization of COVID-19 patient cohorts and assembly of plasma proteomic datasets

Initially, we harmonized patient stratification for COVID-19 severity based on WHOscore (“Severe” vs “NonSevere”) across the three Olink proteomic datasets. In particular, COVID-19 cases who had a fatal outcome or were admitted in the ICU or were intubated were designated as “severe” and the residual cases were designated as “non-severe”. In the MGH study, we designated 80 severe and 225 non-severe cases. In the Mayo study, we demarcated 268 severe and 181 non-severe COVID-19 cases. Furthermore, we determined 24 severe and 40 non-severe cases in the Stanford study. Associations with respective WHOscores and age can be seen in (Sup. Figure 1).

From all 3 Olink studies, 1463 common plasma proteins were bioinformatically studied within APNet and were used for downstream processing to uncover predictive markers of severity.

### 3.2 Data preprocessing and detection of severity drivers across proteomic studies

Next, we used the NetBID2 toolkit through APNet to detect common differentially active proteins (DAPs) in severe COVID-19 cases, for all three Olink studies. Notably, for MGH, the prominent positive drivers included TACSTD2, BAG3, POLR2F, DPY30, and CAPG. Conversely, the top negative drivers for MGH were CCL22, BTC, IGFBP3, TNFSF11, and ICOSLG (Sup. Figure 2A). Similarly, in the case of Mayo, the leading positive drivers consisted of VSIG4, IL1RL1, IL27, KRT19, and JUN, while negative drivers entailed CDON, CD1C, ITGB7, TNFSF11, and LRRN1 (Sup. Figure 2B). For Stanford, the primary positive drivers were LGALS1, CSTB, MAD1L1, DDAH1, and CCL7 and the negative were EPCAM, CPA2, CDNF, DSG4, and CD1C. Pathway enrichment showed that these severe COVID-19 top-drivers were associated with cell migration, monocyte activation, methylation changes and immune cell dysregulation (Sup. Figure 2A-C).

From hereon, we focused on the commonly perturbed drivers across the three studies. APNet captured 333 common differentially active proteins (DAPs) across the three studies and encompassed 163 differentially expressed proteins (DEPs) and 170 hidden drivers (i.e., hidden in at least one of the three Olink datasets) (Figure 3A-B). Among the 333 common drivers, 150 were differentially hyper-active and 183 were hypo-active in severe COVID-19. When analyzing the STRINGdb network of common DAPs, centrality analysis prioritized DEPs like immuno-regulatory interleukins IL4-IL6, keratin modulators (KRT19), chemokines for macrophages and neutrophils (CXCL8/CCL20) and transcription factors (JUN). Other central decreased DEPs were effectors of T cell activation and proliferation (CD8A, CD28), mediators of developmental pathways like the SCF/c-Kit pathway (KIT/KITLG/IL7R/FLT3/CD34) and cellular adhesion surface molecules (ITGB1). Similarly, central hyper-active hidden drivers (“positive”) pertained to growth factors (HGF), ECM remodellers (metalloprotease inhibitor TIMP1), chemoattractants of monocytes, natural killer and T-cells (CXCL9/CXCL10/CCL3) and biomarkers of systemic organ failure (the lipocalin LCN2 indicating acute kidney injury). Other central hypo-active hidden drivers included cellular adhesion molecules (NCAM1, ITGB2, ITGAV), growth factors (FLT3LG, ligand for the FLT3 receptor found in DEPs) or cognate receptors (EGFR, receptor for the Epidermal Growth Factor) and the tumour suppressor molecule PTEN (Figure 3C).

**Figure 3.**
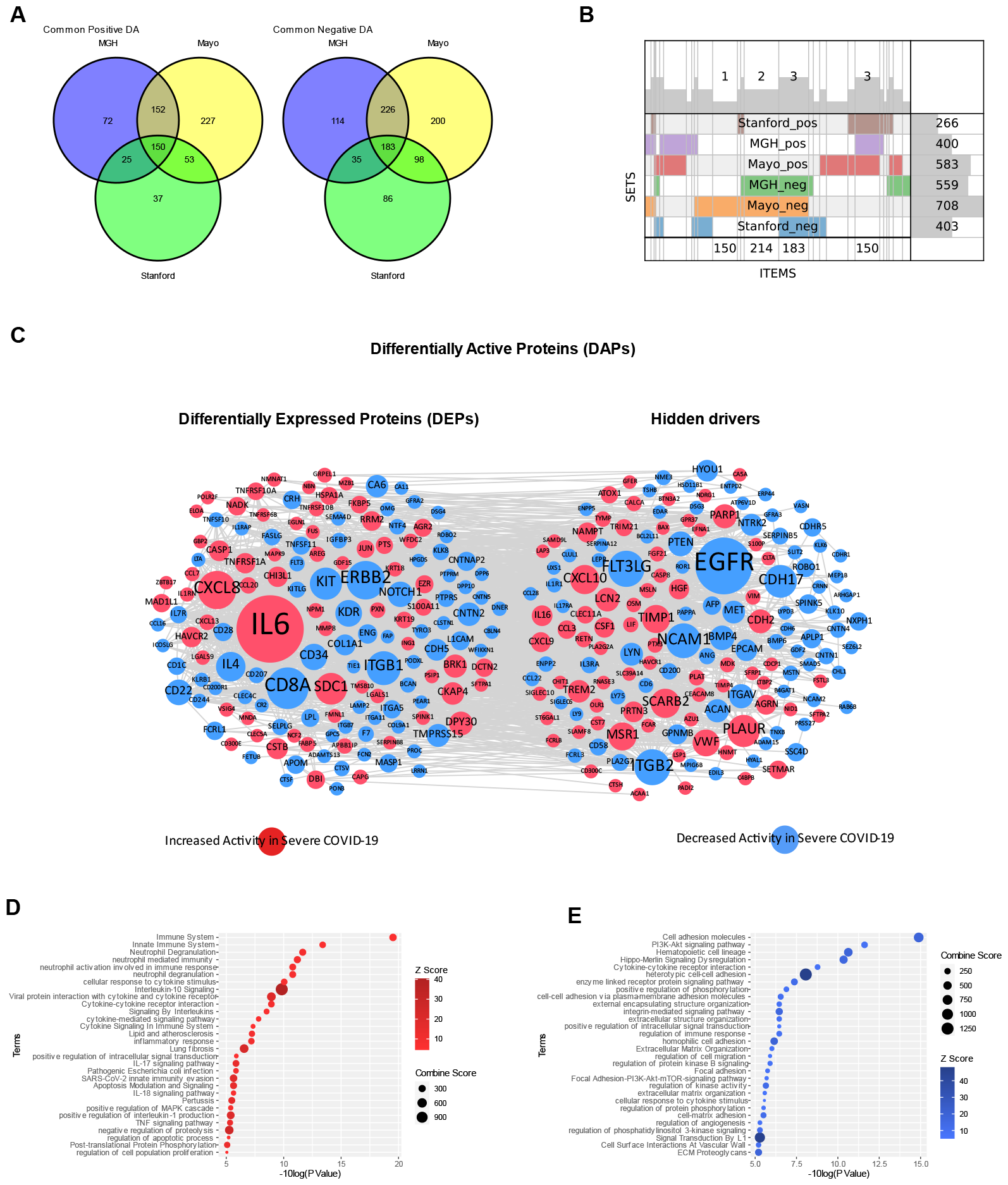
APNet uncovers a large COVID-19 perturbational proteomic space underpinning the 3 distinct Olink datasets (MGH, Mayo, Stanford). **(A)** Venn diagrams showing overlapping differentially expressed proteins (DEPs) and differentially active proteins (DAGs) among the three studies. **(B)** SuperVenn diagram depicting the joint differentially expressed (increased/decreased) or active proteins (hyper/hypo-active) in severe COVID-19 compared to non-severe COVID-19 cases, across the three Olink studies. **(C)** STRINGdb protein-protein interaction networks for joint DEPs and hidden drivers of severity across the three studies (STRINGdb score > 0.4). The size of the nodes is analogous to the centrality of each protein/driver (BetweenessCentrality algorithm) and the colour denotes perturbational direction (red for increased, blue for decreased). **(D-E)** Bubble plots depicting over-representation analysis based on the Enrichr Knowledge Graph (Wikipathways 2021, Reactome, GO:BP, KEGG) for joint drivers with increased (D) and decreased activity (E) in severe COVID-19 cases, among the three Olink plasma proteomic studies.

Pathway enrichment through the Enrichr KG (KEGG, Wikipathways, Reactome, GO:BP) highlighted several biological ground truths involved in COVID-19 immunopathology such as increased activation of innate immunity, lung fibrosis, MAPK signaling, Sars-CoV-2 immuno-evasion, neutrophil degranulation and viral protein interaction with cytokines and cognate receptors (Figure 3D). Conversely, dwindling pathways in severe COVID-19 included the hematopoietic system, inhibitors of the PI3K-Akt signaling pathway, cellular adhesion mechanisms through integrins and the Hippo-Merlin signaling pathway, revealing an impairment of physiological proliferation and migration for circulating immune cells (Figure 3E).

To better dissect the increased perturbational space captured by APNet, distinct cellular enrichment for DEPs and hidden drivers was conducted using the GTEx_Tissues database through Enrichr. DEPs exhibited an over-representation for peripheral blood, spleen, liver, brain, and adipose tissue. Hidden drivers, conversely, implicated other organs like oesophagus, tibial nerve and the cardio-vascular system (Sup. Figure 3A-D). Ensuing pathway enrichment with WikiPathways and GO:BP uncovered an expected affiliation of DEPs with key COVID-19 molecular “landmarks” like apoptosis, viral life cycle, neutrophil degranulation, PI3K Akt signaling and impediments in synapse functionality and angiogenesis. Interestingly, the hidden drivers were skewed towards aberrant insulin signaling, cellular adhesion imbalances (L1cam interactions), propagation of hypoxia and abnormal neuronal behaviour (increased neuroinflammation, decreased neuroplasticity) (Sup. Figure 3E-F, Sup. Figure 4).

These preliminary findings underline the importance of employing activity transformations on distinct COVID-19 plasma proteomic datasets using APNet. Beyond mere differential expression, this approach identified shared, systemic damages caused by Sars-CoV-2 across multiple organs and tissues (Supplementary Material 2-3).

### 3.3 APNet classifies severe COVID-19 cases among distinct plasma proteomic studies

At this point, we hypothesized that the newly discovered hidden drivers had untapped biological potential, which could enhance the clinical prediction of severe COVID-19 cases from plasma proteomics.

We used the common DAPs across the three Olink studies for the ensuing clinical predictions. After initial training in the MGH dataset (for details, see Materials and Methods), APNet accurately predicted severe COVID-19 patients during the early testing phase (MGH-Mayo experiment) with significant robustness (AUC = 0.96, F1 score = 0.9). Biological explainability highlighted the prognostic significance of various DEPs (JUN, IL6, MAPK9, TNFRSF1A, AREG, NTF4, NCF2, TNFRSF10A, FLT3, CKAP4, FLT3LG, SDC1, TNFRSF10B, TNFSF11) but also several hidden drivers (FTL3LG, LYN, PTEN, EFNA1, ACAA1, HGF, TIMP1) (top-20). The most predictive pathways involved the ground-truth “cytokine storm”, MAPK signaling, vascular damage reflected on atherosclerosis potential, protein folding (through HSP90 chaperone), and PI3K-Akt signaling pathway (Figure 4A-B).

**Figure 4.**
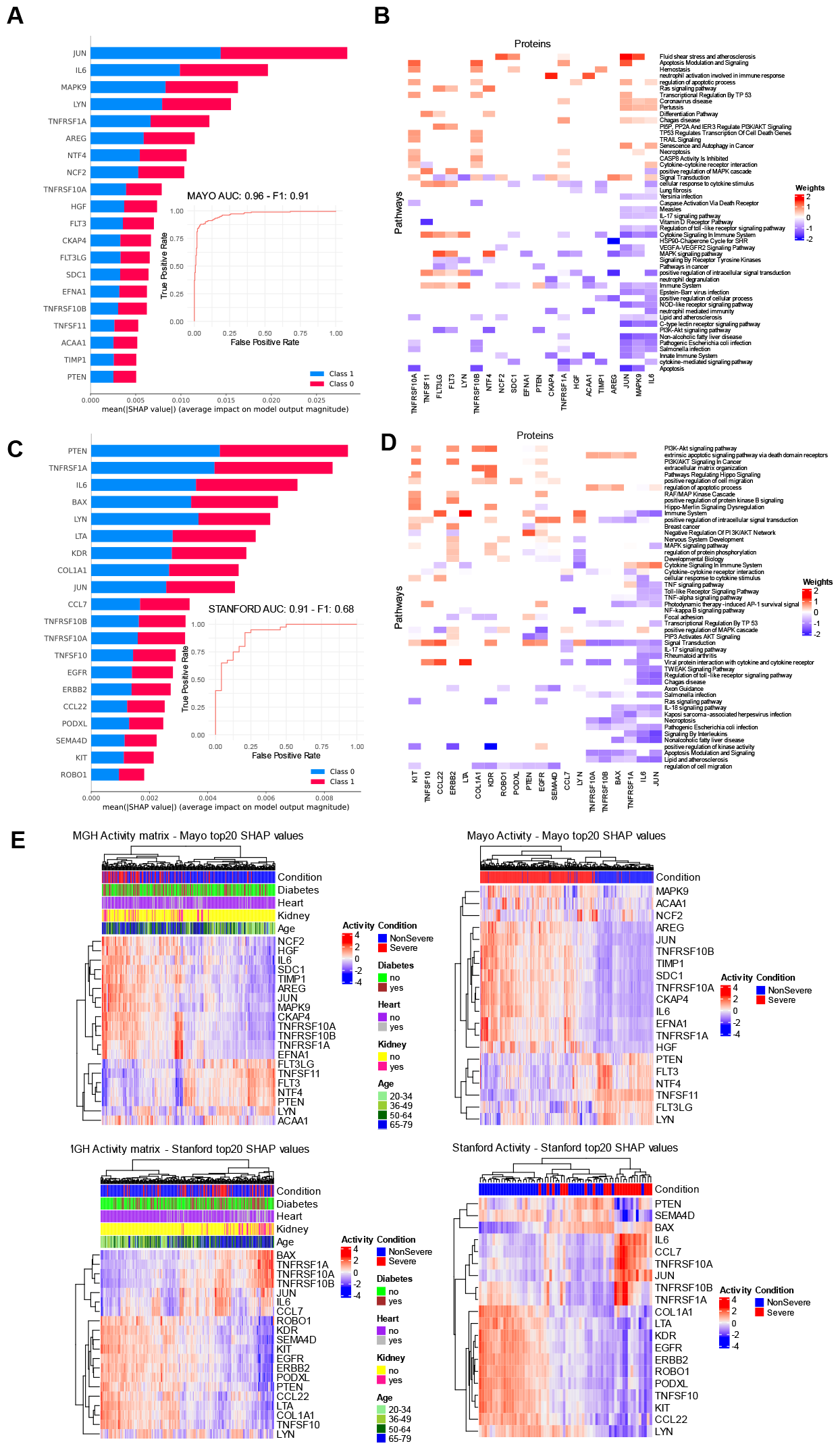
APNet deploys sparse regularization of driver-pathway connections through the PASNet Deep Learning model and robustly classifies severe from non-severe COVID-19 cases in the three Olink proteomic studies. **(A-B**) Bar plots for SHAP values and driver-pathway mapping from PASNet signifying the top-20 predictive drivers and their corresponding pathways, for the MGH (training) / Mayo (testing) experiment. Furthermore, the AUC and F1-score values are depicted. (**C-D**) Same as (A) for the MGH (training) / Stanford (testing) experiment. Class 0 refers to nonsevere and Class 1 refers to severe COVID-19 cases. (**E**) Hierarchical clustering of MGH, Mayo and Stanford cases on the basis on the predictive proteomic drivers, along with selected clinical covariates.

During the second testing phase (MGH-Stanford experiment), APNet once again exhibited significant predictive robustness since, on the Stanford dataset, it could foreshadow severe COVID-19 efficiently (AUC = 0.91, F1 score = 0.68). Biological explainability revealed predictive drivers of severity, many of which overlapped with the ones from the previous testing experiment (i.e. PTEN, JUN, IL6, LYN, TNFRSF1A, TNFRSF10A, TNFRSF10B, TNFSF10) but also unveiled novel ones (BAX, LTA, KDR, COL1A1, CCL7, EGFR, ERBB2, CCL22, PODXL, SEMA4D, KIT, ROBO1). The hidden drivers were PTEN, BAX, CCL22, EGFR, LYN, ROBO1 (Figure 4C).

The most predictive pathways in this experiment involved viral infection and disruption of cytokines and cognate receptors, PI3K-Akt signaling pathway, MAPK signaling, Hippo-Merlin signaling dysregulation, and intensified Interleukin signaling pathway, apoptotic TRAIL signaling, neurotoxicity concerning axonogenesis, and imbalances in lipid metabolism (Figure 4D) (Supplementary Material 4).

Hierarchical clustering on all cases across studies revealed associations of the most predictive drivers with COVID-19 severity, while in the MGH study, further associations with diabetes and kidney disease were also uncovered (Figure 4E).

### 3.4 APNet bridges plasma proteomics with single-cell transcriptomics

At this point, we decided to use APNet for a joint analysis between bulk plasma proteomics (MGH dataset) and scRNA-seq data from circulating peripheral blood mononuclear cells (PBMCs, 4 severe and 10 non-severe MGH cases). We sought to (a) prioritize which predictive drivers of COVID-19 severity could be important for both-omic modalities and (b) trace the cellular origin of various predictive drivers of COVID-19 severity from all the insofar classification experiments among the PBMC cellular populations. (Figure 5A).

**Figure 5.**
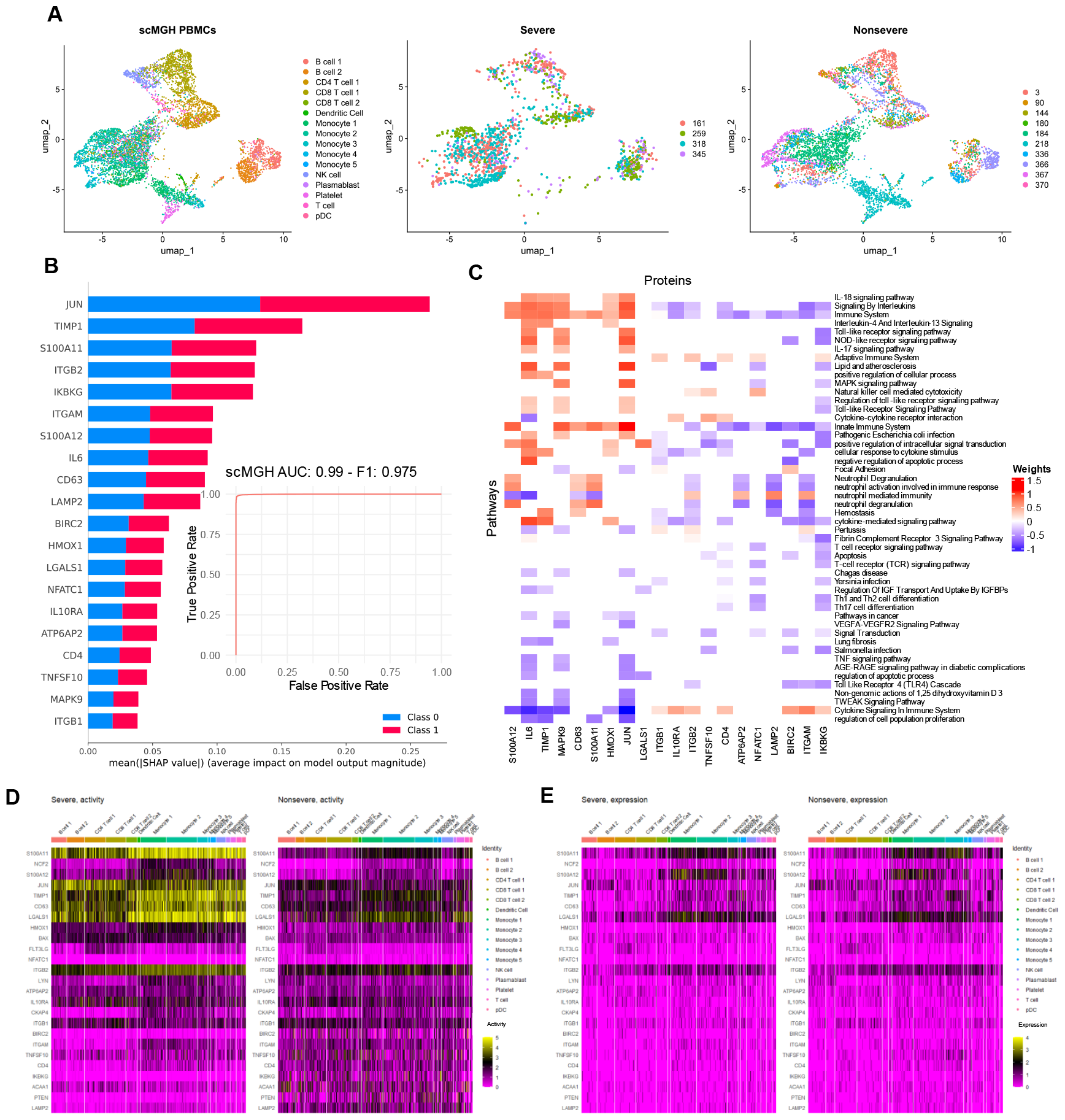
APNet classifies severe COVID-19 cases across multi-omic studies. (A) scRNA-seq data from the Villani group for 14 MGH cases. **(B-C)** Bar plots for SHAP values and driver-pathway mapping from PASNet signifying the top-20 predictive drivers and their corresponding pathways, for the MGH plasma proteomic (training) / scRNA-seq (testing) experiment. Furthermore, the AUC and F1-score values are depicted. Class 0 refers to nonsevere and Class 1 refers to severe COVID-19 cases. **(D-E)** Heatmaps depicting the activity and the expression of predictive drivers from all the APNet experiments, across various PBMC cell types. The scMINER toolkit and visualisation performed the activity calculations were attained through Seurat. Only the predictive drivers with positive activity values are depicted.

Initially, we deployed the scMINER toolkit to convert the typical sparse scRNA-seq expression matrix into a non-sparse activity matrix based on the SJARACNe/MICA/MINIE algorithms (see Materials and Methods for details). Single-cell differential activity analysis revealed 282 differentially active drivers (140 DEGs and 142 hidden drivers) in severe COVID-19, which were also perturbed in the MGH plasma proteomic analysis (Sup. Figure 5A). STRINGdb PPI network modeling and pathway enrichment implicated several key COVID-19 severity drivers in innate/adaptive immunity, viral replication, inflammatory signaling, cell adhesion and lipid metabolism (e.g., *IL6, NCAM, LYN, PTEN, ITGB1, ITGAM*) (Sup. Figure 5B-D), in line with our findings from the previous plasma proteomic analyses.

Next, we trained APNet on the MGH plasma proteomic dataset and tested it on the MGH single-cell dataset (scMGH). APNet was highly robust in classifying severe COVID-19 cases (AUC: 0.99, F1-score: 0.975). The driver-pathway heatmaps pointed towards expected inflammatory and immune pathways (e.g., IL18 signaling, TLR4 stimulation, T cell differentiation) as predictive signalling motifs of severe COVID-19. Five predictive drivers from the previous plasma proteomic experiments were found as predictive genes *(MAPK9, TIMP1, JUN, IL6, TNFSF10)*. The other multi-omic predictive drivers were *S100A12, CD63, LAMP2, BIRC2, HMOX1, LGALS1, NFATC1, IL10RA, ATP6AP2, CD4, ITGB1* (Figure 5B-C).

Lastly, we probed for the single-cell activity profile of various predictive drivers from the MGH-Mayo/MGH-Stanford/MGH-scMGH experiments. We discovered that the most active drivers in severe cases were *JUN* (B/T cells) and *TIMP1* (all PBMCs except B cells and NKs). In contrast, in non-severe cases, it was *PTEN* (monocytes and platelets) and *ACAA1* (all PBMCs but especially B cells). Like *ACAA1*, which opposed its proteomic counterpart, *CKAP4* was also increased in non-severe cases (monocytes). Other active genes in all non-severe PBMCs were *FLT3LG, BAX, LYN* and *TNFSF10*, while *NCF2* was mainly in severe monocytes (Figure 5D-E) (Supplementary Material 5).

Overall, these results elaborate on the cellular origins of certain predictive drivers for severe COVID-19 inferred by APNet in PBMCs and were attained through APNet’ noticeable versatility in bridging across-omic modalities.

### 3.5 Benchmarking APNet against alternative ML/DL methods

To benchmark APNet’s significant performance on COVID-19 classification tasks, we initially retrieved from the literature the predictive models published by the authors of the MGH study (Filbin et al.), the Stanford study (Feyarts et al.), and of an independent study from Qatar which used the MGH study for independent validation. As shown in Table 1, APNet outperformed the MGH and Stanford models (Table 1). Although APNet showed similar performance to Qatar’s predictive model (AUC > 0.95, training-testing on the authors’ in-house data) in demarcating severe COVID-19 cases, it outperformed Qatar’s model in terms of generalizability. This was evident as the latter achieved an AUC of 0.79 when independently tested on the MGH study (Table 1).

**Table 1.** Published ML/DL analyzing MGH and Stanford Olink datasets.

| Study | AI model | AUC |
| --- | --- | --- |
| Filbin et al. (MGH study) | Random Forest (elastic-net logistic regression with cross-validation) | 0.85 |
| Fayerts et al. (Stanford study) | (LASSO) linear regression | 0.77 - 0.79 (Stanford study) |
| Al-Nesf et al. (Qatar study with Boruta algorithm) | MUVR | >0.959 (Qatar data)<br>0.76 (D0) (MGH validation) |

At this point, we performed more specific benchmarking experiments using (a) a variation of APNet where we provided only DEPs to the DL model instead of DAPs (PASNet-expression) and (b) an alternative variation where we substituted the PASNet architecture with one of the most widely used, explainable Machine Learning approaches like Random Forest (RF). The training, validation and testing datasets remained the same as before.

Noticeably, APNet outperformed all alternative DL/ML models based on activity or expression data regarding AUC and F1-score (Figure 6A - B). More specifically, the PASNet-expression model performed poorly on the Mayo dataset (AUC: 0.645, F1 score: 0.7475), and none of the predicted molecules were hidden drivers. Biological explainability indicated ground-truth biological pathways related to COVID-19 immunopathology were the most predictive pathways. However, some more nuanced pathways that APNet retrieved during the MGH-Mayo experiment were missing or under-represented (e.g., lipid imbalances) (Figure 6C-D).

**Figure 6.**
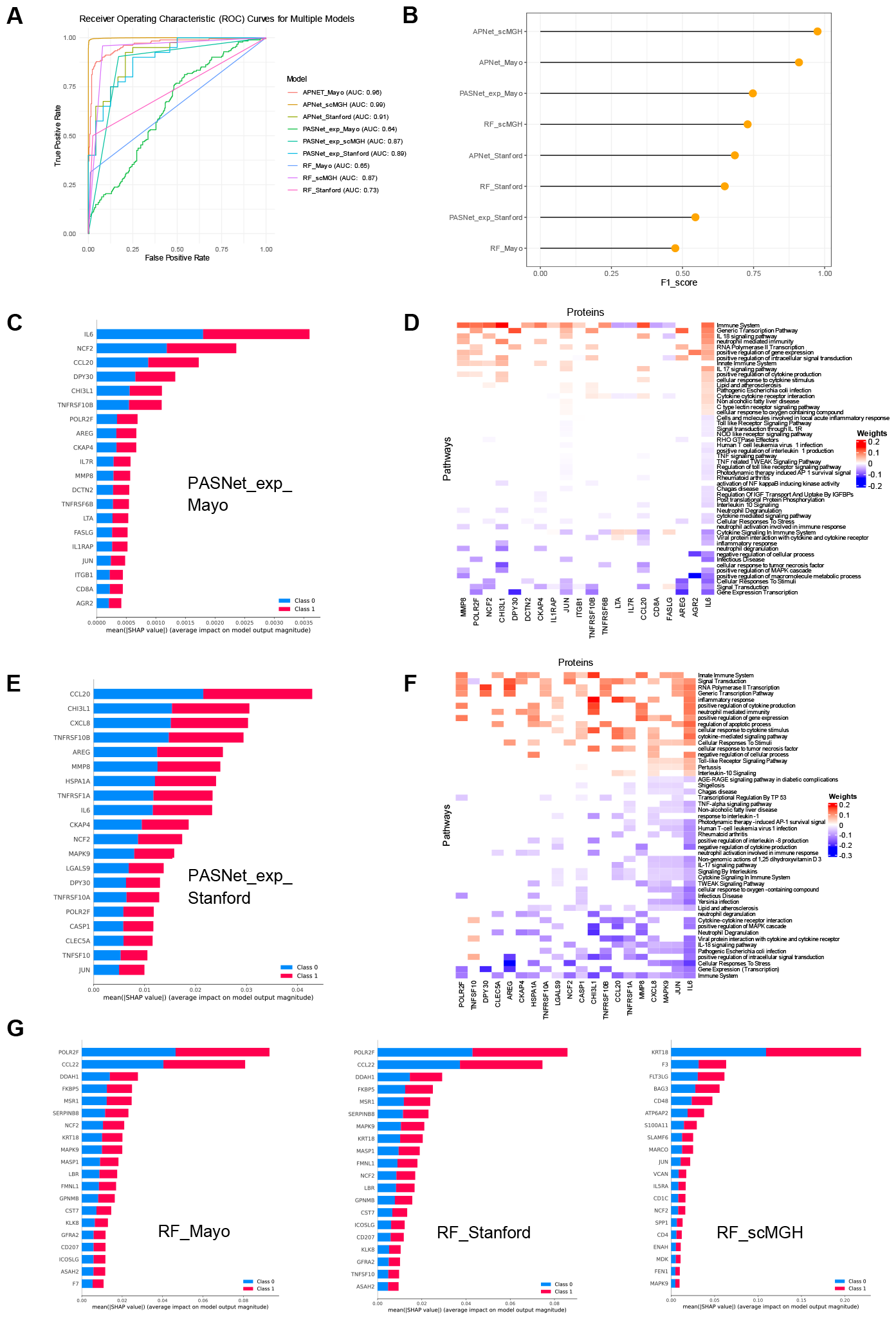
APNet outperforms alternative ML/DL models in classifying severe and non-severe COVID-19 cases. **(A)** ROC curves depicting the performance of APNet and alternative approaches in classifying severe from non-severe cases in respective experiments. Distinct AUC scores are referenced also. **(B)** Lollipop plot depicting the F1-scores from each model from (A). **(C-F)** Barplots with SHAP values for the top 20 most predictive plasma proteins of COVID-19 severity and protein-pathway mapping from PASNet-expression model for MGH-Mayo experiment (C-D) and MGH-Stanford experiment (E-F). **(G)** Barplots with SHAP values showing top 20 most predictive drivers of severity based on the Random Forest (RF) alternative model for the various classification experiments. Class 0 refers to non-severe and Class 1 refers to severe COVID-19 cases.

The PASNet-expression model also under-performed in the Standford study compared to APNet, with an AUC of 0.89 and an F1-score of 0.54. Unsurprisingly, this expression-driven investigation in the Stanford study could only reveal a limited scope of predictive biological pathways like Cellular response to stress, positive regulation of intracellular signaling transduction, Neutrophil degradation, and viral protein response (Figure 6E-F).

The second alternative model based on Random Forest (RF) under-performed even more on Mayo and Stanford datasets than the previous one since the models were validated with AUC: 0.65, F1-score: 0.4746, and AUC: 0.7375, F1-score: 0.6486, respectively, for each dataset. Noticeably, the top-predictive proteins were almost identical across the Mayo and Stanford datasets analysis. Concerning the multi-omic experiment, we opted not to test the PASNet expression-driven model. This decision was based on the intrinsic sparsity of the scRNA-seq data’s expression and the apparent requirement for specific data harmonization or more advanced ML/DL manipulations, which were beyond the scope of our current project. Consequently, we exclusively employed the RF model on the shared perturbational space identified through activity analysis between plasma proteomics and scRNA-seq data. This approach underperformed compared to APNet, as evidenced by an AUC of 0.87 and an F1-score of 0.73 (Figure 6G).

With regards to associations with clinico-biological covariates, the most predictive proteins or drivers from the benchmarking studies exhibited correlations with COVID-19 severity but not to the extend that APNet’s results did (e.g., this is evident in the expression-PASNet MGH-Mayo/Stanford and the RF MGH-Stanford experiments). Furthermore, associations with diabetes and kidney disease were not as straightforward as in the case of APNet (Supplementary Figure 6-7) (Supplementary Material 6).

### 3.6 *In silico* evaluation of APNet’s results based on COVID-19 curated prior knowledge

To evaluate the degree of COVID-19 ground truths that APNet and the other classification models recovered, we mapped each model’s top 20 most predictive proteins from the various experiments to the SIGNOR 3.0 COVID-19 Hallmark pathways (i.e. *Virus Entry, Cytokine storm, Inflammation, Fibrosis, Apoptosis, Innate response to dsRNA, MAPK Activation, ER stress and Stress granules, https://signor.uniroma2.it/covid/*). APNet’s most predictive drivers from the MGH-Mayo and the multi-omic experiments were considerably over-represented (1.5 to 2 fold) on the SIGNOR 3.0 COVID-19 Hallmark pathways than their counterparts from the PASNet-expression and RF models. Concerning the MGH-Stanford experiment, APNet and PASNet-expression exhibited almost an equal number of mappings but different in type, while the RF model was again significantly under-represented. In the case of MAPK activation, a cardinal pathway in COVID-19 pathobiology, APNet accomplished approximately twice as many mappings (8) as the expression-PASNet model (4), revealing higher robustness in connecting predictive drivers of severity with COVID-19 biological underpinnings (Table 2) (Supplementary material 7).

**Table 2.** Biological benchmarking of APNet vs PASNet Expression and RF-Activity. The table measures the number of top-20 predictive drivers that were mapped to the respective SIGNOR 3.0 pathway networks, in each classification experiment (SIGNOR 3.0 COVID-19 Hallmarks).

|  | MGH-Mayo |  |  | MGH-Stanford |  |  | scMGH |  |
| --- | --- | --- | --- | --- | --- | --- | --- | --- |
| **SIGNOR 3.0 COVID-19 Hallmark** | **APNet** | **PASNet-Expression** | **RF-Activity** | **APNet** | **PASNet-Expression** | **RF-Activity** | **APNet** | **RF-Activity** |
| *Virus Entry* | 4 | 2 | 1 | 4 | 5 | 1 | 4 | 2 |
| *Cytokine Storm* | 5 | 5 | 0 | 5 | 7 | 0 | 7 | 3 |
| *Inflammation* | 6 | 5 | 0 | 9 | 7 | 0 | 7 | 2 |
| *Fibrosis* | 4 | 3 | 1 | 3 | 5 | 1 | 7 | 4 |
| *Apoptosis* | 9 | 5 | 1 | 8 | 7 | 1 | 5 | 2 |
| *Innate response to dsRNA* | 3 | 3 | 0 | 3 | 5 | 0 | 4 | 2 |
| *MAPK activation* | 6 | 3 | 1 | 8 | 5 | 1 | 6 | 3 |
| *ER stress* | 3 | 2 | 0 | 4 | 3 | 0 | 3 | 1 |
| *Stress granules* | 5 | 3 | 1 | 5 | 6 | 1 | 6 | 3 |
| *Total mappings* | 45 | 31 | 5 | 49 | 50 | 5 | 49 | 22 |

### 3.7 APNet enables the creation of weighted graph models for mechanistic hypotheses: The case of ACAA1

We postulated that combining SJARACNe co-expression networks, with pathways that APNet ingested as biological priors before classification tasks and the weights it assigned to them upon completion of demarcating severe COVID-19 could be helpful to *in silico* predict regulatory motifs and signaling patterns driving severe COVID-19.

To demonstrate this feature, we focused on the MGH-Mayo experiment, we assembled a multipartite graph of with driver-driver and driver-pathway connections and we sought to leverage information about ACAA1 (Acetyl-CoA Acyltransferase 1), which was one of the top 20 most predictive drivers, was designated as a hidden driver by our analysis and it was significantly hypo-active in severe PBMCs which suggested the its plasma proteomic signature derived from an alternative tissue or organ.

By retrieving the MGH SJARACNe SHAP graph (a series of small positive, coherent feedforward loops with NCF2, TIMP1, CKAP4 as sources, TNFRSF10B as a significant “sink” and FLT3 as the primary inhibitor), it was apparent that no obvious connection existed between ACAA1 and the other predictive drivers (Figure 7A). To validate the biological plausibility of the SJARACNe graph, the respective PPI network from STRINGdb was leveraged (interaction score > 0.4), indicating high interconnectivity for most of the predictive drivers. Interestingly, ACAA1 and CKAP4 remained unconnected (https://version-12-0.string-db.org/cgi/network?networkId=bmlgZrzN1Cex) (Figure 7B).

**Figure 7.**
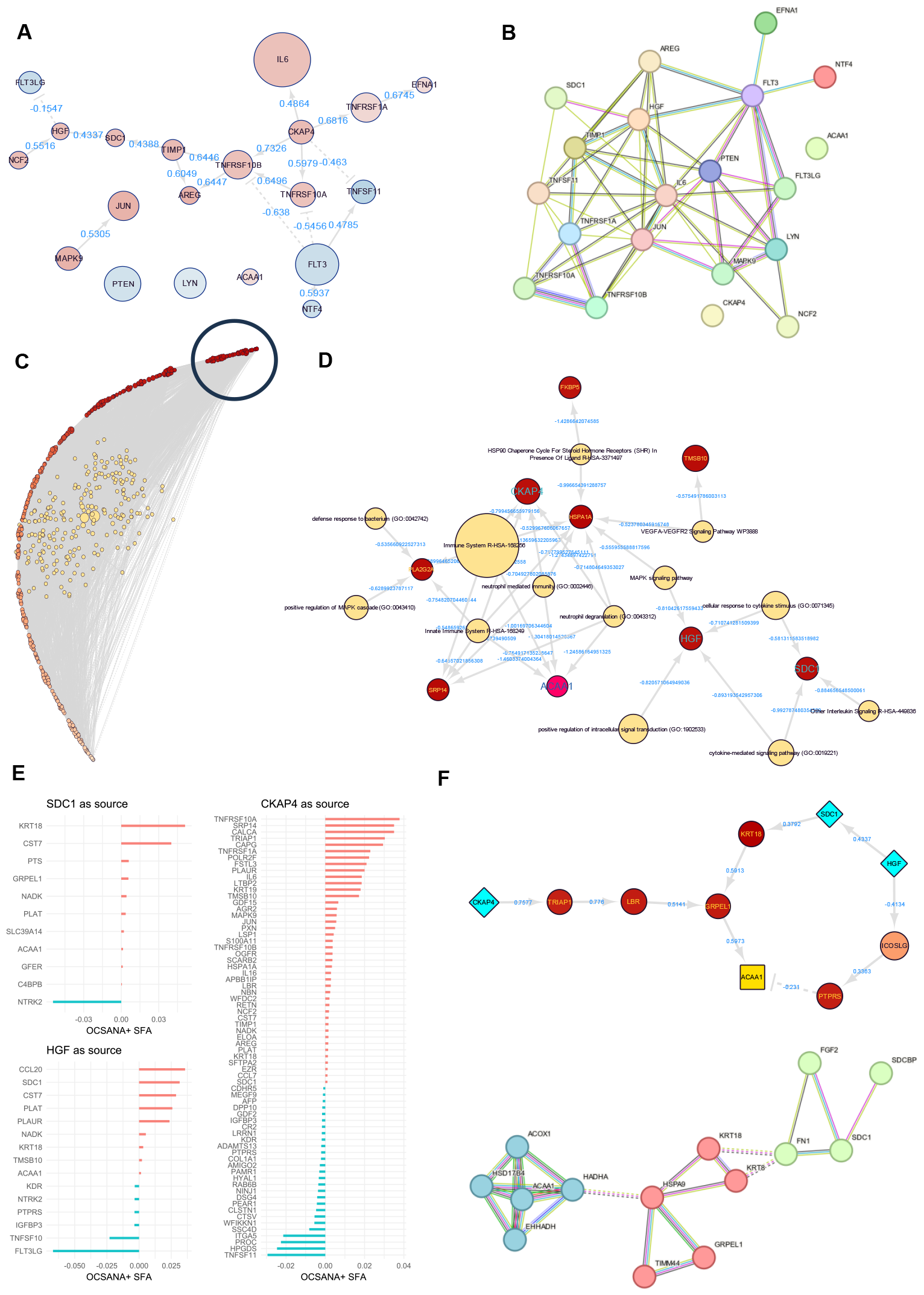
APNet enables the assembly of complex graphs that can be leveraged to discover non-apparent connections of ACAA1 with other predictive drivers of COVID-19 severity. **(A)** SJARACNe co-expression (adj.pvalue<0.05 for MI calculation) directed network from APNet’s complex graph showing the interactions of the top 20 most predictive drivers of COVID-19 severity for the MGH-Mayo scenario. **(B)** STRINGdb network with (interaction score > 0.4) for the same severity drivers. **(C)** APNet’s complex graph after tSNE dimensionality reduction using the clusterMaker app in Cytoscape, based on the “liver” score from the TISSUES 2.0 database for each driver/node. The darkest colour denotes a higher liver-specific association. The most liver-specific cluster of drivers is designated within the circle **(D)** Part of APNet’s complex graph showing highly liver-specific drivers and connected pathways with high prognostic significance, based on their PASNet weights. **(E)** Lollipop plots showing the SFA scores from the OCSANA+ app in Cytoscape in each node, after signal propagation from SDC1, CKAP4 and HGF on the entire APNet complex graph. **(F)** Part of APNet’s complex graph showing shortest paths based on the PathLinker app in Cytoscape starting from CKAP4, SDC1 and HGF and extending towards ACAA1. Below, the STRINGdb equivalent PPI network (interaction score > 0.4) of the SDC1-KRT18-GRPEL1-ACAA1 path with intermediate nodes provided by STRINGdb, after k-means clustering.

Next, considering that ACAA1 is predominantly expressed in the liver based on our previous GTEx analysis, we took inspiration from representation learning (Zitnik et al., 2019) and performed dimensionality reduction on the APNet complex graph with the tSNE algorithm, looking for maximum variance in liver expression based on TISSUES 2.0 scores. A distinct cluster with highly liver-specific drivers was detected. To gain a better insight on them, we isolated their subgraph with their most prognostic connected pathways (PASNet weight > 0.5 and < -0.5). We detected a graph “island” which contained four highly predictive drivers of COVID-19 severity among other proteins (ACAA1, SDC1, HGF, CKAP4) and connected pathways involved Immune System signaling, neutrophil degranulation, MAPK signaling, chaperone activation (HSP90) and VEGF signaling (Figure 7C-D).

Based on these findings, we posited that there should be an underlying connection between ACAA1 and some of the other three predictive drivers of severity. We resorted to the OCSANA+ Cytoscape application to simulate signal propagation from SDC1, HGF and CKAP4 on the APNet complex graph. By calculating the Signal Flow Analysis (SFA, see Materials and Methods) metric, it became apparent that HGF and SDC1 signal propagation converged towards ACAA1 through various intermediate proteins. A similar effect on ACAA1 was not observed in the case of CKAP4, which did not appear to propagate any signal towards ACAA1 (Figure 7E).

To better elucidate these findings, we calculated the shortest paths from SDC1, HGF and CKAP4 towards ACAA1 using the PathwayLinker application on Cytoscape. When selecting the “additive weight method” for the MI score as edge weight, PathLinker highlighted 2 critical shortest paths: (a) a signaling cascade commencing from SDC1 and reaching ACAA1 through KRT18 and GRPEL1 and (b) an incoherent feed-forward loop starting from HGF and through inhibiting ICOSLG which activated PTPRS which inhibited ACAA1. The “unweighted method” in PathwayLinker returned the same results. Notwithstanding, when selecting the “probabilistic weight method” for the MI score as edge weight, PathwayLinker suggested a larger signaling cascade commencing from CKAP4, and extending through TRIAP1, LBR and GRPEL1 towards ACAA1 (Figure 7F).

To computationally validate these APNet shortest paths, we queried the STRINGdb database for the respective PPI networks. After 2 rounds of expansion, we retrieved a singular PPI network (14 nodes, 23 edges, https://version-12-0.string-db.org/cgi/network?networkId=b57tJ84II6T8), connecting SDC1, GRPEL1, KRT18 and ACAA1 which upon k-means clustering revealed three components relative to fatty-acid metabolism (ACAA1, ACOX1, HADHA, HSD17B4, EHHADH), mitochondrial protein transport (HSPA9, KRT18, KRT8, GRPEL1, TIMM44) and cell surface interactions (FN1, SDC1, SDCBP, FGF2). In the case of the other shortest paths, the corresponding STRINGdb queries required more than 5 cycles of expansion to produce PPI networks encompassing all drivers of interest (CKAP4: 55 nodes, 375 edges, https://version-12-0.string-db.org/cgi/network?networkId=bjbqEWsYzPw4; HGF: 44 nodes, 214 edges, https://version-12-0.string-db.org/cgi/network?networkId=bcT9lsyOCwJ7) (Sup. Figure 8).

Finally, as an additional step to assess the potential significance of these paths in a more COVID-19-specific biological context, we queried the BYCOVID19 data portal (https://www.covid19dataportal.org/) for the “COVID-19 association score” provided by the OpenTargets platform. SDC1 exhibited the highest score (0.555) with a considerable difference from some of the other drivers of severity (KRT18=0.05, LBR=0.004, CKAP4=0.006, HGF=0.025), confirming the biological prioritization of the SDC1-ACAA1 nascent connection that APNet uncovered (Supplementary Material 8-9).

## 4. Discussion - Conclusion

In the current work, focusing on COVID-19 omics, we present APNet, a computational DL pipeline to elucidate complex biological motifs while classifying patients based on their clinical severity.

APNet is inspired by computational approaches modeling Gene Regulatory Networks (GRNs), which have been instrumental in discovering new interactions between biological entities and formulating novel scientific hypotheses. APNet combines some of the best practices in the field by combining an Information Theory model (SJARACNe algorithm) through a Bayesian scope (NetBID2/scMINER toolkits) (Delgado and Gómez-Vela, 2019) and a biologically-informed neural network with enhanced explainability (PASNet and SHAP values) for supervised patient clustering. The above bioinformatic tools have been shown independently to effectively discover potential biomarkers and druggable targets in diseases however, to the best of our knowledge, they have never been used as a unified pipeline for COVID-19 or any other disease type (Wang et al., 2021) (Ding et al., 2023) (Hao et al., 2018).

In our study, we utilized APNet to predict severe COVID-19 cases in three different Olink plasma proteomic datasets (MGH, Mayo, Stanford), and a complementary scRNA-seq study. APNet conducted biologically informed predictions using driver-pathway associations (KEGG, Reactome, GO:BP, Wikipathways) with remarkable robustness, outperforming alternative ML/DL approaches which either lacked (a) the activity transformations enabled by the NetBID2/scMINER toolkits (PASNet-expression model) or (b) the PASNet DL architecture (Random Forest classifications). Based on the biological explainability of each model (SHAP values, driver-pathway mapping with learning DL weights) and COVID-19 curated biological ground-truths (SINGOR COVID-19 pathway networks), evidently, APNet was able to better approximate the systemic nature of severe COVID-19 from the provided biological data. We posit that APNet performed so efficiently due to the sparse regularization of the hierarchical relationships of drivers and pathways after initial differential activity analysis. Hence, APNet was able to capture both well known but also more nuanced perturbations in severe COVID-19 (i.e., known drivers but also “hidden drivers” like ACAA1, FLT3) implicating several potential tissues of origin and a diverse repertoire of critical pathways. Indicatively, some of the most predictive drivers and pathways that APNet captured concerned apoptosis, dishevelled PI3K-Akt stimulation (FLT3/FLT3LG, PTEN, NTF4, KIT), neurodegeneration (EGFR, SEMAD4), cell differentiation (TNFSF11), neutrophil degranulation (ACAA1), lipid metabolism (TNFRSF10A), immune and interleukin signaling (CD63, TIMP1, JUN), T cell receptor signaling (BIRC2, NFATC1, CD4, IKBKG), oxidative phosphorylation (ATP6AP2). These signaling cascades and some of these drivers have already been implicated with COVID-19, which attests to APNet’s overall capacity to make biologically plausible predictions. (Basile et al., 2022; Chidambaram et al., 2022; Merad and Martin, 2020; Pistollato et al., 2022; Thompson et al., 2021). A paradigmatic case concerning the translational value of APNet’s findings was the implication of MAPK pathway in severe COVID-19 based on various drivers (e.g., MAPK9, AREG, KIT, JUN, FLT3LG). These drivers were not prioritized to the same extend as highly predictive by the alternative ML/DL models – if prioritized at all. This could explain in part why APNet surpassed these models as a classifier of COVID-19 severity since components of the MAPK pathway (sH-RAS, C-RAF, MAPK1, MAPK2 and ERK) have emerged as critical tenets of Sars-CoV-2 tropism in PBMCs and have been associated with adverse clinical covariates like hypoxia, dyspnoea and vascular damages (Cusato et al., 2023).

Finally, APNet extends beyond biological explainability to actionability regarding the formulation of mechanistic hypotheses, by providing the capacity to generate a weighted driver-pathway network that incorporates information from SJARACNe co-expression networks, the differential activity analysis, the PASNet DL clinical predictions and external dedicated bioinformatic databases like STRINGdb. APNet enabled through graph representation learning, shortest path detection, and signal propagation simulation the prediction of a liver-specific signaling cascade in severe COVID-19 involving ACAA1 (*hidden driver with prognostic significance but no apparent connections to other predictive drivers*), SDC1, KRT18, and GRPEL1. These predictions are not biologically implausible given the implication of SDC1 and KRT18 in inflammation and epithelial damage (Ghondaghsaz et al., 2023; Liao et al., 2020), the involvement of the mitochondrial GRPEL1 in host/Sars-CoV-2 interactions (Zhang et al., 2022), the clinical correlation of ACAA1 (a mediator of fatty acid oxidation in the mitochondria and the peroxisomes) with ICU-admittance in COVID-19 (Penrice-Randal et al., 2022) and a severe mitochondria dysfunction in the liver of severe COVID-19 cases (Guarnieri et al., 2023).

The work herein is not without its limitations. One limitation concerns the restricted number of studies involved and the binary assignment of drivers to pathways. Pathway activation is a dynamic process controlled by fluctuations in expression or activity changes of a protein or drivers, respectively. Outputs from more advanced pathway enrichment techniques like GSEA could be more instructive for the DL model to perform classifications more aptly. Another limitation is the need to perform several manual steps in APNet’s complex graphs to test hypotheses and leverage new insights, which might hinder data exploration and analysis. Another issue worth noting is that APNet does not include clinical covariates as clinical-biological priors, which could be addressed in the future by adopting in our pipeline the more clinically-oriented version of Cox-PASNet (Hao et al., 2019).

Overall, APNet is a robust pipeline that can simplify the extraction of intricate biological insights from complex biological data while also performing clinical predictions and testing mechanistic hypotheses. *In vitro/in vivo* validations should accompany future implementations of APNet to validate the pipeline’s true translational credibility. Additionally, APNet’s scalability to other multi-factorial disease-omic datasets (such as cancer and neurodegenerative diseases) should be explored along with its potential deployment in other computational tasks (like multi-omic data integration and interactions with knowledge graph pipelines).

## Supporting information

Supplemental material for online view

## Abbreviations

APNet: Activity PASNet
ARDS: Acute Respiratory Distress Syndrome
AUC: Area Under the Curve
DAPs: Differential Active Proteins
DEPs: Differential Expressed Proteins
DL: Deep Learning
DOME: Data Optimization, Model, Evaluation
EXP: Expression Values
ICU: Intensive Care Unit
KG: Knowledge Graph
Mayo: Mayo Clinic
MGH: Massachusetts General Hospital
MI: Mutual Information
MLP: Multi-Layer Perceptron
NPX: Normalized Protein eXpression
PASNet: Pathway-Associated Sparse Deep Neural Network
PBMC: Peripheral Blood Mononuclear Cell
PEA: Proximity Extension Assay
RF: Random Forest
ROC: Receiver Operating Characteristic
SFA: Signal Flow Analysis
SHAP: Shapley additive explanation
Stanford: Stanford Hospital
SVM: Support Vector Machine
WHO: World Health Organization
XAI: eXplainable Artificial Intelligent

## Disclosures

None.

## Funding

This work was funded by the HORIZON-INFRA-2021-EOSC-01-04 project **Scilake** (https://scilake.eu/). Also, this work was also supported by ELIXIR (https://elixir-europe.org/), the research infrastructure for life-science data.

## Competing interests

None.

## Author Contributions

Conceptualization, resources, methodology, investigation, formal analysis, writing – original draft, writing – review & editing: G.I.G, V.I.V., S.D. Supplementary analysis by G.K. Supervision: A.G., G.A.P., F.P.

## Data Availability

APNet R and Python scripts and the code to re-create the figures of this manuscript can be found at https://github.com/BiodataAnalysisGroup/APNet.

The datasets used in this study can be accessed in the following links: ***MGH Olink proteomics***: https://info.olink.com/mgh-covid-study-overview-page?hsCtaTracking=fff99a2a-81c1-4e4a-a70d-6922d26503b4%7C202c2809-0976-48f7-aad0-3903c36624ca, ***Mayo Olink Proteomics***: https://www.thelancet.com/journals/landig/article/PIIS2589-7500(22)00112-1/fulltext#supplementaryMaterial, ***Stanford Olink proteomics***: https://datadryad.org/stash/dataset/doi:10.5061/dryad.9cnp5hqmn, ***single-cell MGH Villani group***: https://www.covid19cellatlas.org/index.patient.html. All of the above data are also included in our Zenodo link: 10.5281/zenodo.10438830

Other supplementary material can be found at the “Supplementary Materials for Online” folder and on Zenodo: 10.5281/zenodo.10438830

## Description of supplementary materials

| File name | Description |
| --- | --- |
| Supplementary_Material_1_DOME.pdf | DOME recommendations for the APNet models |
| Supplementary_Material_2_MGH_Mayo_Stanford_matrices.xlsx | Expression, and Activity matrices for MGH, Mayo, and Stanford datasets |
| Supplementary_Material_3_DA_MGH_Mayo_Stanford.xlsx | Differential Activity outputs of MGH, Mayo, Stanford and there relevant master file |
| Supplementary_Material_4_APNet_weights.xlsx | APNet activity s1_weights outputs for MGH-Mayo, MGH-Stanford |
| Supplementary_Material_5_scMGH_MGH.xlsx | Differential Activity outputs of scMGH + APNet activity s1_weights outputs for MGH-scMGH |
| Supplementary_Material_6_PASNet_Expr_Benchmarking.xlsx | PASNet expression s1_weights outputs for MGH-Mayo, MGH-Stanford |
| Supplementary_Material_7_SIGNOR_COVID_Hallmarks.cys | Cytoscape file that contains the SIGNOR 3.0 COVID-19 hallmark pathways and the mapping of the most predictive drivers from APNet and the benchmarking experiments |
| Supplementary_Material_8_The_ACAA1_case_study.cys | Cytoscape file that contains the bipartite driver-pathway weighted network based on the MGH SJARACNe co-expression graph (MGH-Mayo experiment). This bipartite network is used to leverage information about ACAA1. |
| Supplementary_Material_9_The_ACAA1_case_study_OCSANA.txt | Txt file guiding through the Cytoscape analysis for the ACAA1 case study. |

## Notes

https://zenodo.org/records/10438830

https://github.com/BiodataAnalysisGroup/APNet

### Competing Interest Statement

The authors have declared no competing interest.

## Bibliography

Babačić, H., Christ, W., Araújo, J.E., Mermelekas, G., Sharma, N., Tynell, J., García, M., Varnaite, R., Asgeirsson, H., Glans, H., Lehtiö, J., Gredmark-Russ, S., Klingström, J., Pernemalm, M., 2023. Comprehensive proteomics and meta-analysis of COVID-19 host response. Nat. Commun. 14, 5921. 10.1038/s41467-023-41159-z

Basile, M.S., Cavalli, E., McCubrey, J., Hernández-Bello, J., Muñoz-Valle, J.F., Fagone, P., Nicoletti, F., 2022. The PI3K/Akt/mTOR pathway: A potential pharmacological target in COVID-19. Drug Discov. Today 27, 848–856. 10.1016/j.drudis.2021.11.002

Byeon, S.K., Madugundu, A.K., Garapati, K., Ramarajan, M.G., Saraswat, M., Kumar-M, P., Hughes, T., Shah, R., Patnaik, M.M., Chia, N., Ashrafzadeh-Kian, S., Yao, J.D., Pritt, B.S., Cattaneo, R., Salama, M.E., Zenka, R.M., Kipp, B.R., Grebe, S.K.G., Singh, R.J., Sadighi Akha, A.A., Algeciras-Schimnich, A., Dasari, S., Olson, J.E., Walsh, J.R., Venkatakrishnan, A.J., Jenkinson, G., O’Horo, J.C., Badley, A.D., Pandey, A., 2022. Development of a multiomics model for identification of predictive biomarkers for COVID-19 severity: a retrospective cohort study. Lancet Digit. Health 4, e632–e645. 10.1016/S2589-7500(22)00112-1

Chidambaram, V., Kumar, A., Majella, M.G., Seth, B., Sivakumar, R.K., Voruganti, D., Bavineni, M., Baghal, A., Gates, K., Kumari, A., Al’Aref, S.J., Galiatsatos, P., Karakousis, P.C., Mehta, J.L., 2022. HDL cholesterol levels and susceptibility to COVID-19. eBioMedicine 82, 104166. 10.1016/j.ebiom.2022.104166

Cusato, J., Manca, A., Palermiti, A., Mula, J., Costanzo, M., Antonucci, M., Trunfio, M., Corcione, S., Chiara, F., De Vivo, E.D., Ianniello, A., Ferrara, M., Di Perri, G., De Rosa, F.G., D’Avolio, A., Calcagno, A., 2023. COVID-19: A Possible Contribution of the MAPK Pathway. Biomedicines 11, 1459. 10.3390/biomedicines11051459

Delgado, F.M., Gómez-Vela, F., 2019. Computational methods for Gene Regulatory Networks reconstruction and analysis: A review. Artif. Intell. Med. 95, 133–145. 10.1016/j.artmed.2018.10.006

Diamond, M.S., Kanneganti, T.-D., 2022. Innate immunity: the first line of defense against SARS-CoV-2. Nat. Immunol. 23, 165–176. 10.1038/s41590-021-01091-0

Dimitsaki, S., Gavriilidis, G.I., Dimitriadis, V.K., Natsiavas, P., 2023. Benchmarking of Machine Learning classifiers on plasma proteomic for COVID-19 severity prediction through interpretable artificial intelligence. Artif. Intell. Med. 137, 102490. 10.1016/j.artmed.2023.102490

Ding, L., Shi, H., Qian, C., Burdyshaw, C., Veloso, J.P., Khatamian, A., Pan, Q., Dhungana, Y., Xie, Z., Risch, I., Yang, X., Huang, X., Yan, L., Rusch, M., Brewer, M., Yan, K.-K., Chi, H., Yu, J., 2023. scMINER: a mutual information-based framework for identifying hidden drivers from single-cell omics data. BioRxiv Prepr. Serv. Biol. 10.1101/2023.01.26.523391

Ding, L., Shi, H., Qian, C., Burdyshaw, C., Veloso, J.P., Pan, Q., Dhungana, Y., Xie, Z., Risch, I., Yang, X., Yan, L., Rusch, M., Brewer, M., Yan, K.-K., Chi, H., n.d. scMINER: a mutual information-based framework for identifying hidden drivers from single-cell omics data.

Dong, X., Ding, L., Thrasher, A., Wang, X., Liu, J., Pan, Q., Rash, J., Dhungana, Y., Yang, X., Risch, I., Li, Y., Yan, L., Rusch, M., McLeod, C., Yan, K.-K., Peng, J., Chi, H., Zhang, J., Yu, J., 2023. NetBID2 provides comprehensive hidden driver analysis. Nat. Commun. 14, 2581. 10.1038/s41467-023-38335-6

Eldjarn, G.H., Ferkingstad, E., Lund, S.H., Helgason, H., Magnusson, O.Th., Gunnarsdottir, K., Olafsdottir, T.A., Halldorsson, B.V., Olason, P.I., Zink, F., Gudjonsson, S.A., Sveinbjornsson, G., Magnusson, M.I., Helgason, A., Oddsson, A., Halldorsson, G.H., Magnusson, M.K., Saevarsdottir, S., Eiriksdottir, T., Masson, G., Stefansson, H., Jonsdottir, I., Holm, H., Rafnar, T., Melsted, P., Saemundsdottir, J., Norddahl, G.L., Thorleifsson, G., Ulfarsson, M.O., Gudbjartsson, D.F., Thorsteinsdottir, U., Sulem, P., Stefansson, K., 2023. Large-scale plasma proteomics comparisons through genetics and disease associations. Nature 622, 348–358. 10.1038/s41586-023-06563-x

Evangelista, J.E., Xie, Z., Marino, G.B., Nguyen, N., Clarke, D.J.B., Ma’ayan, A., 2023. Enrichr-KG: bridging enrichment analysis across multiple libraries. Nucleic Acids Res. 51, W168–W179. 10.1093/nar/gkad393

Feyaerts, D., Hédou, J., Gillard, J., Chen, H., Tsai, E.S., Peterson, L.S., Ando, K., Manohar, M., Do, E., Dhondalay, G.K.R., Fitzpatrick, J., Artandi, M., Chang, I., Snow, T.T., Chinthrajah, R.S., Warren, C.M., Wittman, R., Meyerowitz, J.G., Ganio, E.A., Stelzer, I.A., Han, X., Verdonk, F., Gaudillière, D.K., Mukherjee, N., Tsai, A.S., Rumer, K.K., Jacobsen, D.R., Bjornson-Hooper, Z.B., Jiang, S., Saavedra, S.F., Valdés Ferrer, S.I., Kelly, J.D., Furman, D., Aghaeepour, N., Angst, M.S., Boyd, S.D., Pinsky, B.A., Nolan, G.P., Nadeau, K.C., Gaudillière, B., McIlwain, D.R., 2022. Integrated plasma proteomic and single-cell immune signaling network signatures demarcate mild, moderate, and severe COVID-19. Cell Rep. Med. 3, 100680. 10.1016/j.xcrm.2022.100680

Filbin, M.R., Mehta, A., Schneider, A.M., Kays, K.R., Guess, J.R., Gentili, M., Fenyves, B.G., Charland, N.C., Gonye, A.L.K., Gushterova, I., Khanna, H.K., LaSalle, T.J., Lavin-Parsons, K.M., Lilley, B.M., Lodenstein, C.L., Manakongtreecheep, K., Margolin, J.D., McKaig, B.N., Rojas-Lopez, M., Russo, B.C., Sharma, N., Tantivit, J., Thomas, M.F., Gerszten, R.E., Heimberg, G.S., Hoover, P.J., Lieb, D.J., Lin, B., Ngo, D., Pelka, K., Reyes, M., Smillie, C.S., Waghray, A., Wood, T.E., Zajac, A.S., Jennings, L.L., Grundberg, I., Bhattacharyya, R.P., Parry, B.A., Villani, A.-C., Sade-Feldman, M., Hacohen, N., Goldberg, M.B., 2021. Longitudinal proteomic analysis of severe COVID-19 reveals survival-associated signatures, tissue-specific cell death, and cell-cell interactions. Cell Rep. Med. 2, 100287. 10.1016/j.xcrm.2021.100287

Ghondaghsaz, E., Khalaji, A., Norouzi, M., Fraser, D.D., Alilou, S., Behnoush, A.H., 2023. The utility of syndecan-1 circulating levels as a biomarker in patients with previous or active COVID-19: a systematic review and meta-analysis. BMC Infect. Dis. 23, 510. 10.1186/s12879-023-08473-9

Gil, D.P., Law, J.N., Murali, T.M., 2017. The PathLinker app: Connect the dots in protein interaction networks. F1000Research 6, 58. 10.12688/f1000research.9909.1

Guarnieri, J.W., Dybas, J.M., Fazelinia, H., Kim, M.S., Frere, J., Zhang, Y., Soto Albrecht, Y., Murdock, D.G., Angelin, A., Singh, L.N., Weiss, S.L., Best, S.M., Lott, M.T., Zhang, S., Cope, H., Zaksas, V., Saravia-Butler, A., Meydan, C., Foox, J., Mozsary, C., Bram, Y., Kidane, Y., Priebe, W., Emmett, M.R., Meller, R., Demharter, S., Stentoft-Hansen, V., Salvatore, M., Galeano, D., Enguita, F.J., Grabham, P., Trovao, N.S., Singh, U., Haltom, J., Heise, M.T., Moorman, N.J., Baxter, V.K., Madden, E.A., Taft-Benz, S.A., Anderson, E.J., Sanders, W.A., Dickmander, R.J., Baylin, S.B., Wurtele, E.S., Moraes-Vieira, P.M., Taylor, D., Mason, C.E., Schisler, J.C., Schwartz, R.E., Beheshti, A., Wallace, D.C., 2023. Core mitochondrial genes are down-regulated during SARS-CoV-2 infection of rodent and human hosts. Sci. Transl. Med. 15, eabq1533. 10.1126/scitranslmed.abq1533

Hao, J., Kim, Y., Kim, T.-K., Kang, M., 2018. PASNet: pathway-associated sparse deep neural network for prognosis prediction from high-throughput data. BMC Bioinformatics 19, 510. 10.1186/s12859-018-2500-z

Hao, J., Kim, Y., Mallavarapu, T., Oh, J.H., Kang, M., 2019. Interpretable deep neural network for cancer survival analysis by integrating genomic and clinical data. BMC Med. Genomics 12, 189. 10.1186/s12920-019-0624-2

Liang, X., Sun, R., Wang, J., Zhou, K., Li, J., Chen, S., Lyu, M., Li, S., Xue, Z., Shi, Y., Xie, Y., Zhang, Q., Yi, X., Pan, J., Wang, D., Xu, J., Zhu, H., Zhu, G., Zhu, J., Zhu, Y., Zheng, Y., Shen, B., Guo, T., 2023. Proteomics Investigation of Diverse Serological Patterns in COVID-19. Mol. Cell. Proteomics 22, 100493. 10.1016/j.mcpro.2023.100493

Liao, M., Liu, Y., Yuan, J., Wen, Y., Xu, G., Zhao, J., Cheng, L., Li, J., Wang, X., Wang, F., Liu, L., Amit, I., Zhang, S., Zhang, Z., 2020. Single-cell landscape of bronchoalveolar immune cells in patients with COVID-19. Nat. Med. 26, 842–844. 10.1038/s41591-020-0901-9

Marazzi, L., Gainer-Dewar, A., Vera-Licona, P., 2020. OCSANA+: optimal control and simulation of signaling networks from network analysis. Bioinformatics 36, 4960–4962. 10.1093/bioinformatics/btaa625

Merad, M., Martin, J.C., 2020. Pathological inflammation in patients with COVID-19: a key role for monocytes and macrophages. Nat. Rev. Immunol. 20, 355–362. 10.1038/s41577-020-0331-4

Paul, S.G., Saha, A., Biswas, A.A., Zulfiker, Md.S., Arefin, M.S., Rahman, Md.M., Reza, A.W., 2023. Combating Covid-19 using machine learning and deep learning: Applications, challenges, and future perspectives. Array 17, 100271. 10.1016/j.array.2022.100271

Penrice-Randal, R., Dong, X., Shapanis, A.G., Gardner, A., Harding, N., Legebeke, J., Lord, J., Vallejo, A.F., Poole, S., Brendish, N.J., Hartley, C., Williams, A.P., Wheway, G., Polak, M.E., Strazzeri, F., Schofield, J.P.R., Skipp, P.J., Hiscox, J.A., Clark, T.W., Baralle, D., 2022. Blood gene expression predicts intensive care unit admission in hospitalised patients with COVID-19. Front. Immunol. 13, 988685. 10.3389/fimmu.2022.988685

Pistollato, F., Petrillo, M., Clerbaux, L.-A., Leoni, G., Ponti, J., Bogni, A., Brogna, C., Cristoni, S., Sanges, R., Mendoza-de Gyves, E., Fabbri, M., Querci, M., Soares, H., Munoz, A., Whelan, M., Van de Eede, G., 2022. Effects of spike protein and toxin-like peptides found in COVID-19 patients on human 3D neuronal/glial model undergoing differentiation: Possible implications for SARS-CoV-2 impact on brain development. Reprod. Toxicol. 111, 34–48. 10.1016/j.reprotox.2022.04.011

Thompson, E.A., Cascino, K., Ordonez, A.A., Zhou, W., Vaghasia, A., Hamacher-Brady, A., Brady, N.R., Sun, I.-H., Wang, R., Rosenberg, A.Z., Delannoy, M., Rothman, R., Fenstermacher, K., Sauer, L., Shaw-Saliba, K., Bloch, E.M., Redd, A.D., Tobian, A.A.R., Horton, M., Smith, K., Pekosz, A., D’Alessio, F.R., Yegnasubramanian, S., Ji, H., Cox, A.L., Powell, J.D., 2021. Metabolic programs define dysfunctional immune responses in severe COVID-19 patients. Cell Rep. 34, 108863. 10.1016/j.celrep.2021.108863

Walsh, I., Fishman, D., Garcia-Gasulla, D., Titma, T., Pollastri, G., ELIXIR Machine Learning Focus Group, Capriotti, E., Casadio, R., Capella-Gutierrez, S., Cirillo, D., Del Conte, A., Dimopoulos, A.C., Del Angel, V.D., Dopazo, J., Fariselli, P., Fernández, J.M., Huber, F., Kreshuk, A., Lenaerts, T., Martelli, P.L., Navarro, A., Broin, P.Ó., Piñero, J., Piovesan, D., Reczko, M., Ronzano, F., Satagopam, V., Savojardo, C., Spiwok, V., Tangaro, M.A., Tartari, G., Salgado, D., Valencia, A., Zambelli, F., Harrow, J., Psomopoulos, F.E., Tosatto, S.C.E., 2021. DOME: recommendations for supervised machine learning validation in biology. Nat. Methods 18, 1122–1127. 10.1038/s41592-021-01205-4

Wang, F., Morita, K., DiNardo, C.D., Furudate, K., Tanaka, T., Yan, Y., Patel, K.P., MacBeth, K.J., Wu, B., Liu, G., Frattini, M., Matthews, J.A., Little, L.D., Gumbs, C., Song, X., Zhang, J., Thompson, E.J., Kadia, T.M., Garcia-Manero, G., Jabbour, E., Ravandi, F., Bhalla, K.N., Konopleva, M., Kantarjian, H.M., Andrew Futreal, P., Takahashi, K., 2021. Leukemia stemness and co-occurring mutations drive resistance to IDH inhibitors in acute myeloid leukemia. Nat. Commun. 12, 2607. 10.1038/s41467-021-22874-x

Wik, L., Nordberg, N., Broberg, J., Björkesten, J., Assarsson, E., Henriksson, S., Grundberg, I., Pettersson, E., Westerberg, C., Liljeroth, E., Falck, A., Lundberg, M., 2021. Proximity Extension Assay in Combination with Next-Generation Sequencing for High-throughput Proteome-wide Analysis. Mol. Cell. Proteomics 20, 100168. 10.1016/j.mcpro.2021.100168

Zhang, N., Wang, S., Wong, C.C.L., 2022. Proteomics research of SARS-CoV-2 and COVID-19 disease. Med. Rev. 2, 427–445. 10.1515/mr-2022-0016

Zhong, W., Edfors, F., Gummesson, A., Bergström, G., Fagerberg, L., Uhlén, M., 2021. Next generation plasma proteome profiling to monitor health and disease. Nat. Commun. 12, 2493. 10.1038/s41467-021-22767-z

Zitnik, M., Nguyen, F., Wang, B., Leskovec, J., Goldenberg, A., Hoffman, M.M., 2019. Machine learning for integrating data in biology and medicine: Principles, practice, and opportunities. Inf. Fusion 50, 71–91. 10.1016/j.inffus.2018.09.012

