## Supplemental material for online view for "APNet, an explainable sparse deep learning model to discover differentially active drivers of severe COVID-19": Supplementary_Material_1_DOME.pdf

### DOME recommendations table: MGH plasma proteomics-scRNA-seq

| | | |
|---|---|---|
| **DOME** | Version | *1.0* |
| **Data: MGH NPX proteomics** | Provenance | *Olink NPX plasma proteomic COVID-19 from Massachusetts General Hospital (MGH). 223 Proteins.* $N_{covid19}$ *= 293 samples (*$N_{severe}$ *= 76,* $N_{non\text{-}severe}$ *= 216)* |
| | Dataset splits | *All dataset is used as train dataset of the AI model.* |
| | Redundancy between data splits | *No splits* |
| | Availability of data | *Yes, original URL: https://olink.com/application/mgh-covid-19-study/*<br>*URL: HTTPS://GITHUB.COM/BIODATAANALYSISGROUP/APNET*<br>*Free use license.* |
| **Data: MGH scRNAseq** | Provenance | *Single-cells RNAseq COVID-19 from Massachusetts General Hospital (MGH). 223 Genes.* $N_{covid19}$ *= 6665 cells (*$N_{severe}$ *= 1521,* $N_{non\text{-}severe}$ *= 5144)* |
| | Dataset splits | *All dataset is used as validation and test dataset of the AI model.* |
| | Redundancy between data splits | *No splits* |
| | Availability of data | *Yes, original URL: https://www.covid19cellatlas.org/index.patient.html*<br>*URL: HTTPS://GITHUB.COM/BIODATAANALYSISGROUP/APNET*<br>*Free use license.* |
| **Data: Pathways** | Provenance | *Pathway extracted from EnrichR-KG by loading the common critical drivers from Three Studies (MGH, Mayo, Stanford)* |
| | Data splits | *All datasets were used as layer in model* |
| | Redundancy between data splits | *No splits* |
| | Availability of data | *Yes, original URL: https://maayanlab.cloud/enrichr-kg*<br>*URL: HTTPS://GITHUB.COM/BIODATAANALYSISGROUP/APNET*<br>*Free use licence* |
| **Optimization** | Algorithm | *PASNet (Pathway-Associated Sparse Deep Neural Network)*<br>*https://github.com/DataX-JieHao/PASNet*<br>*https://doi.org/10.1186/s12859-018-2500-z* |
| | Meta-predictions | *No* |
| | Data encoding | *1.Activity values as numerical matrices, with samples as rows and the 223 common critical proteins/genes as columns (features) with an addition last column specified the condition of samples/cells. (Same format for 2 datasets)*<br>*2. Binary matrix [0,1] with pathways as row and proteins/genes as column.* |
| | Parameters | *Learning rate = 0.0007 and 0.007 depending on model used. The dropouts for two intermediate layers were also applied with a dropping probability of 0.8 and 0.7, respectively. Adaptive Moment Estimation (Adam) was* |

| | | |
|---|---|---|
| | | *performed as the stochastic optimizer.* |
| | Features | *For Olink MGH plasma proteomics and scRNAseq MGH datasets we have 223 proteins as features.* |
| | Fitting | *PASNet optimizes a small sub-network, which involves feasible nodes and parameters to train instead of the whole network and then makes the sub-network sparse, to avoid overfitting in high dimensional low sample data (HDLS).* |
| | Regularization | *Sparse deep learning architecture represents multiple molecular biological layers which use sparse regularization. L2 regularization = 0.005 and 0.0003 depending on model used.* |
| | Availability of configuration | *URL: HTTPS://GITHUB.COM/BIODATAANALYSISGROUP/APNET*<br>*Free use licence* |
| **Model** | Interpretability | *Black box, as correlation between input and output is masked. The model could be mentioned as partially transparent because of the pathway layer. The explainability method of SHAP values is used as an added explanation method.* |
| | Output | *Classification of patients/cells based on disease severity condition.* |
| | Execution time | *~2h (Depends on Computational Power)* |
| | Availability of software | *URL: HTTPS://GITHUB.COM/BIODATAANALYSISGROUP/APNET*<br>*Free use licence* |
| **Evaluation** | Evaluation method | *Independent datasets* |
| | Performance measures | *F1-score, AUC* |
| | Comparison | *Independent benchmarking with alternative ML/DL models.*<br>*Run PASNet with Expression values of 3 datasets.*<br>*Run Random Forest Classification with Activity values of 3 datasets* |
| | Confidence | *Not applicable.* |
| | Availability of evaluation | *URL: HTTPS://GITHUB.COM/BIODATAANALYSISGROUP/APNET*<br>*Free use licence* |

#### DOME recommendations table: MGH-Mayo-Stanford plasma proteomics

| | | |
|---|---|---|
| **DOME** | Version | *1.0* |
| **Data: MGH NPX proteomics** | Provenance | *Olink NPX plasma proteomic COVID-19 from Massachusetts General Hospital (MGH). 250 Proteins.* $N_{covid19}$ *= 305 samples (*$N_{severe}$ *= 80,* $N_{non\text{-}severe}$ *= 225)* |
| | Dataset splits | *All dataset is used as train dataset of the AI model.* |
| | Redundancy between data splits | *No splits* |
| | Availability of data | *Yes, original URL: https://olink.com/application/mgh-covid-19-study/*<br>*URL: HTTPS://GITHUB.COM/BIODATAANALYSISGROUP/APNET*<br>*Free use license.* |
| **Data: Mayo NPX proteomics** | Provenance | *Olink NPX plasma proteomics COVID-19 from patients visited one of the three Mayo Clinic sites in the USA (Minnesota, Arizona, or Florida). 250 Proteins.* $N_{covid19}$ *= 305 samples (*$N_{severe}$ *= 268,* $N_{non\text{-}severe}$ *= 181)* |
| | Dataset splits | *All dataset is used as validation and test dataset of the AI model.* |
| | Redundancy between data splits | *No splits* |
| | Availability of data | *Yes, original URL: https://doi.org/10.1016/S2589-7500(22)00112-1, appendix*<br>*URL: HTTPS://GITHUB.COM/BIODATAANALYSISGROUP/APNET*<br>*Free use license.* |
| **Data: Stanford NPX proteomics** | Provenance | *Olink NPX plasma proteomics COVID-19 from Stanford Medicine. 250 Proteins.* $N_{covid19}$ *= 64 samples (*$N_{severe}$ *= 24,* $N_{non\text{-}severe}$ *= 40)* |
| | Dataset splits | *All dataset is used as test dataset of the AI model.* |
| | Redundancy between data splits | *No splits* |
| | Availability of data | *Yes, original URL: https://datadryad.org/stash/dataset/doi:10.5061/dryad.9cnp5hqmn*<br>*URL: HTTPS://GITHUB.COM/BIODATAANALYSISGROUP/APNET*<br>*Free use license.* |
| **Data: Pathways** | Provenance | *Pathway extracted from EnrichR-KG by loading the common critical drivers from Three Studies (MGH, Mayo, Stanford)* |
| | Data splits | *All datasets were used as layer in model* |
| | Redundancy between data splits | *No splits* |
| | Availability of data | *Yes, original URL: https://maayanlab.cloud/enrichr-kg*<br>*URL: HTTPS://GITHUB.COM/BIODATAANALYSISGROUP/APNET* |

| | | |
|---|---|---|
| | | *Free use licence* |
| **Optimization** | Algorithm | *PASNet (Pathway-Associated Sparse Deep Neural Network)*<br>*https://github.com/DataX-JieHao/PASNet*<br>*https://doi.org/10.1186/s12859-018-2500-z* |
| | Meta-predictions | *No* |
| | Data encoding | *1.Activity values as numerical matrices, with samples as rows and the 250 common critical proteins as columns (features) with an addition last column specified the condition of samples. (Same format for 3 datasets)*<br>*2. Binary matrix [0,1] with pathways as row and proteins as column.* |
| | Parameters | *Learning rate = 0.0007 and 0.007 depending on model used. The dropouts for two intermediate layers were also applied with a dropping probability of 0.8 and 0.7, respectively. Adaptive Moment Estimation (Adam) was performed as the stochastic optimizer.* |
| | Features | *For all 3 datasets we have 250 proteins as features.* |
| | Fitting | *PASNet optimizes a small sub-network, which involves feasible nodes and parameters to train instead of the whole network and then makes the sub-network sparse, to avoid overfitting in high dimensional low sample data (HDLS).* |
| | Regularization | *Sparse deep learning architecture represents multiple molecular biological layers which use sparse regularization. L2 regularization = 0.005 and 0.0003 depending on model used.* |
| | Availability of configuration | *URL: HTTPS://GITHUB.COM/BIODATAANALYSISGROUP/APNET*<br>*Free use licence* |
| **Model** | Interpretability | *Black box, as correlation between input and output is masked. The model could be mentioned as partially transparent because of the pathway layer. The explainability method of SHAP values is used as an added explanation method.* |
| | Output | *Classification of patients/cells based on disease severity condition.* |
| | Execution time | *~2h (Depends on Computational Power)* |
| | Availability of software | *URL: HTTPS://GITHUB.COM/BIODATAANALYSISGROUP/APNET*<br>*Free use licence* |
| **Evaluation** | Evaluation method | *Independent datasets* |
| | Performance measures | *F1-score, AUC* |
| | Comparison | *Run Random Forest Classification with Activity values of 2 datasets* |
| | Confidence | *Not applicable.* |
| | Availability of evaluation | *URL: HTTPS://GITHUB.COM/BIODATAANALYSISGROUP/APNET*<br>*Free use licence* |
