## Supplemental material for online view for "APNet, an explainable sparse deep learning model to discover differentially active drivers of severe COVID-19": Supplementary_Material_9_The_ACAA1_case_study_guide.txt

The ACAA1 case study
--------------------
The purpose of this case study is to uncover obscure connections of ACAA1, a hidden and highly predictive driver of severe COVID-19 based on the MGH-Mayo experiment with other highly predictive drivers.
To do so, we are going to analyze the driver-pathway bipartite graph based on the MGH dataset. The analysis is performed in Cytoscape and the apps used are "String", "Pathlinker" and "OCSANA+".

1. Perform tSNE in the MGH bipartite graph (MGH-Mayo experiment) based on the TISSUES "liver" node attribute and obtain the liver cluster (the most liver-specific cluster).

2. Retrieve the most predictive pathways affecting the liver and the liver drivers: 
	- Select adjacent edges --> Filter >0.5, <-0.5 --> New network with selected edges

3. Detect possible association of ACAA1 with CKAP4, HGF, SDC1 due to driver-pathway network

4. Then, return to the MGH bipartite graph and use Shortest Paths to uncover hidden connections among CKAP4, HGF, SDC1

5. Firstly, choose only drivers (~330 nodes) and then, using Pathlinker, MI as weight, and 2 different algorithms to approach the paths, find overlaps:
	- "weights are additive"
	- "weights are probabilities" 

Edge weights are additive: The score of a path is the
sum of the weights of the edges in it. PathLinker com-
putes the k paths of lowest score in this case as well.


Edge weights are probabilities: This situation arises
often with protein interactions networks, since such a
weight indicates the experimental reliability of an edge.
PathLinker treats the edge weights as multiplicative
and computes the k highest cost paths, where the cost
of a path is the product of the edge weights. Internally,
PathLinker transforms each weight to the absolute value of
its logarithm to map the problem to the additive case.

Isolate potential shortest paths. Potential connections through intermediate nodes appear to be formed among ACAA1 and SDC1, HGF and CKAP4.


6. To further prioritize the abovementioned shortest paths, OCSANA+ can be used to simulate signal propagation:
By analyzing the MGH driver network, select OCSANA app and "Signal Flow Analysis" and select as activated nodes iteratively SDC1, HGF and CKAP4

Signal Flow Analysis (SFA):
SDC1>>>>>
KRT19: 0.0
IL6: 0.0
PLAT: 0.022821773229381923
ADGRG2: 0.0
ACAA1: 0.019304425594079203
PTPRS: 0.0
CKAP4: 0.0
FSTL3: 0.0
DSG4: 0.0
CALCA: 0.0
NCF2: 0.0
ICOSLG: 0.0
FAP: 0.0
HGF: 0.0
LTBP2: 0.0
PTS: 0.05103103630798288
POLR2F: 0.0
CXCL13: 0.0
TNFSF10: 0.0
CTSV: 0.0
ITGA11: 0.0
GRPEL1: 0.054601160977938205
SRP14: 0.0
TNFRSF10A: 0.0
TNFRSF10B: 0.0
CST7: 0.10206207261596575
KRT18: 0.17677669529663687
COL1A1: 0.0
IL7R: 0.0
NADK: 0.03605086100812361
SDC1: 0.5
APBB1IP: 0.0
CCL20: 0.0
IGFBP3: 0.0


CKAP4>>>>>>>
KRT19: 0.07095369570699615
IL6: 0.07241681243618579
PLAT: 0.012527444621948387
ADGRG2: -0.011240298573294415
ACAA1: 0.012902596427527419
PTPRS: -0.03227522974103959
CKAP4: 0.5
FSTL3: 0.0721017220162472
DSG4: -0.0443112618421153
CALCA: 0.08333333333333333
NCF2: 0.025515518153991435
ICOSLG: -0.02696818384160967
FAP: -0.01692741742761445
HGF: 0.0
LTBP2: 0.08333333333333333
PTS: 0.0
POLR2F: 0.07593591053487325
CXCL13: 0.02248059714658883
TNFSF10: -0.019593976524998197
CTSV: -0.02248059714658883
ITGA11: -0.019917582994779477
GRPEL1: 0.004218823974232251
SRP14: 0.08333333333333333
TNFRSF10A: 0.08333333333333333
TNFRSF10B: 0.0829270707215016
CST7: 0.008534403190923549
KRT18: 0.0
COL1A1: -0.018633899812498245
IL7R: -0.006588078458684123
NADK: 0.014614434943119821
SDC1: 0.0
APBB1IP: 0.024056261216234408
CCL20: 0.010452466538471627
IGFBP3: -0.036299758273496706


HGF>>>>>>>
KRT19: 0.0
IL6: 0.0
PLAT: 0.07002577590769957
ADGRG2: 0.0
ACAA1: 0.011852021146094311
PTPRS: -0.022307953230637757
CKAP4: 0.0
FSTL3: 0.0
DSG4: 0.0
CALCA: 0.0
NCF2: 0.0
ICOSLG: -0.04759718605670352
FAP: 0.0
HGF: 0.5
LTBP2: 0.0
PTS: 0.0
POLR2F: 0.0
CXCL13: 0.0
TNFSF10: -0.06454972243679029
CTSV: 0.0
ITGA11: 0.0
GRPEL1: 0.011214624862040824
SRP14: 0.0
TNFRSF10A: 0.0
TNFRSF10B: 0.0
CST7: 0.06454972243679029
KRT18: 0.0
COL1A1: 0.0
IL7R: 0.0
NADK: 0.03884860009775963
SDC1: 0.0
APBB1IP: 0.0
CCL20: 0.07905694150420947
IGFBP3: -0.010206207261596574

Based on the above, we can discern which is the most probable and robust connection of ACAA1 with other potential predictive drivers (in our case SDC1 primarily).
