## Supplementary Figures for "APNet, an explainable sparse deep learning model to discover differentially active drivers of severe COVID-19"

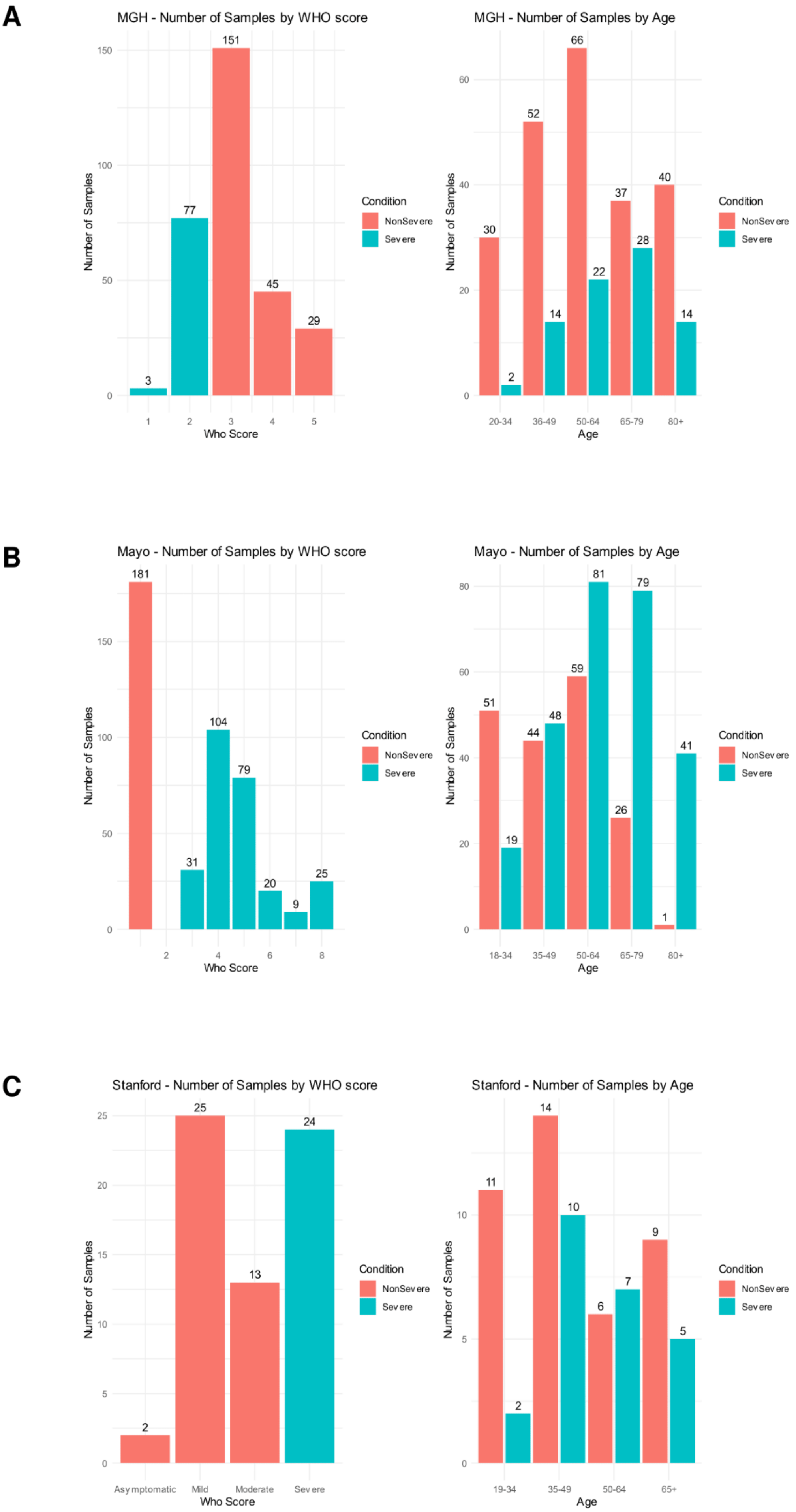
A
MGH - Number of Samples by WHO score
Number of Samples
Who Score
Condition
NonSevere
Severe
3
77
151
45
29
MGH - Number of Samples by Age
Age
20-34
35-49
50-64
65-79
80+
30
2
52
14
66
22
37
28
40
14
B
Mayo - Number of Samples by WHO score
181
31
104
79
20
9
25
Mayo - Number of Samples by Age
18-34
35-49
50-64
65-79
80+
51
19
44
48
59
81
26
79
1
41
C
Stanford - Number of Samples by WHO score
Asymptomatic
Mild
Moderate
Severe
2
25
13
24
Stanford - Number of Samples by Age
19-34
35-49
50-64
65+
11
2
14
10
6
7
9
5

**Supplementary Figure 1: COVID-19 patient stratification based on WHOscore and age across all 3 Olink proteomic studies. (A-C)** Barplots displaying the distribution of samples within specific WHO-score categories for three Olink datasets [MGH (**A**), Mayo (**B**), and Stanford (**C**)] and the corresponding age distribution across severe and non-severe COVID-19 cases.

A

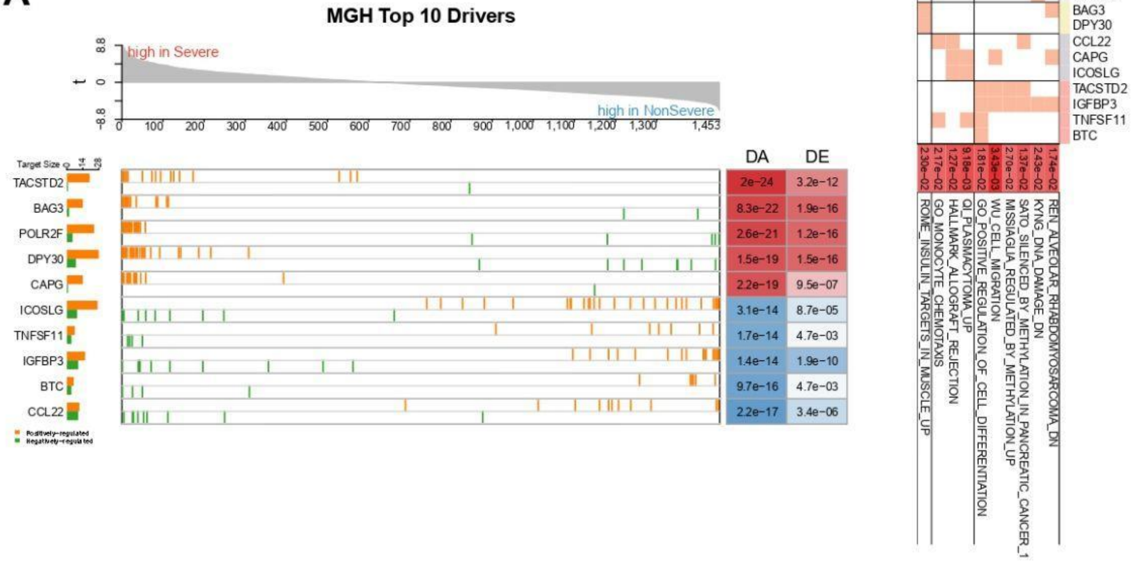
MGH Top 10 Drivers
high in Severe
high in NonSevere
t
Target Size
TACSTD2
BAG3
POLR2F
DPY30
CAPG
ICOSLG
TNFSF11
IGFBP3
BTC
CCL22
Positively-regulated
Negatively-regulated
DA
DE
2e-24
3.2e-12
8.3e-22
1.9e-16
2.6e-21
1.2e-16
1.5e-19
1.5e-16
2.2e-19
9.5e-07
3.1e-14
8.7e-05
1.7e-14
4.7e-03
1.4e-14
1.9e-10
9.7e-16
4.7e-03
2.2e-17
3.4e-06

B

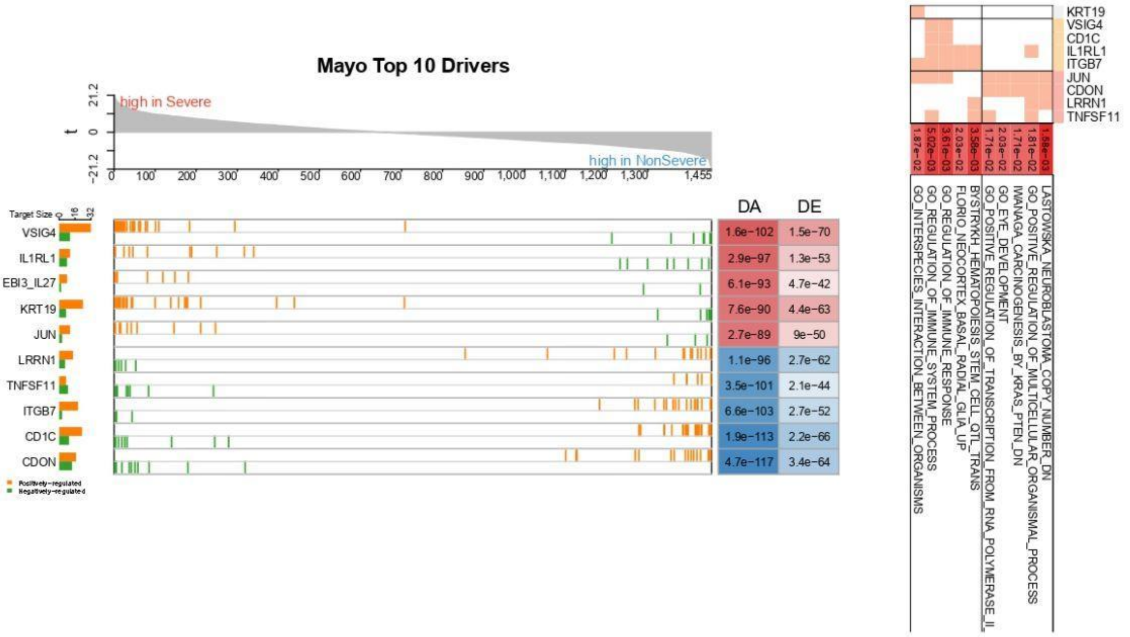
Mayo Top 10 Drivers
high in Severe
high in NonSevere
t
Target Size
VSIG4
IL1RL1
EBI3_IL27
KRT19
JUN
LRRN1
TNFSF11
ITGB7
CD1C
CDON
Positively-regulated
Negatively-regulated
DA
DE
1.6e-102
1.5e-70
2.9e-97
1.3e-53
6.1e-93
4.7e-42
7.6e-90
4.4e-63
2.7e-89
9e-50
1.1e-96
2.7e-62
3.5e-101
2.1e-44
6.6e-103
2.7e-52
1.9e-113
2.2e-66
4.7e-117
3.4e-64

C

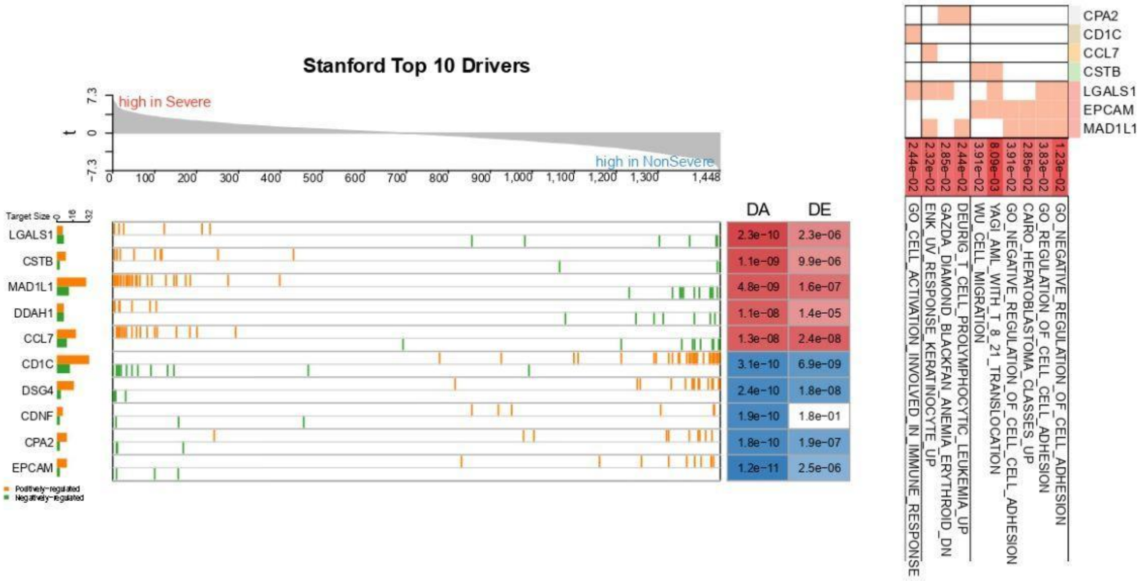
Stanford Top 10 Drivers
high in Severe
high in NonSevere
t
Target Size
LGALS1
CSTB
MAD1L1
DDAH1
CCL7
CD1C
DSG4
CDNF
CPA2
EPCAM
Positively-regulated
Negatively-regulated
DA
DE
2.3e-10
2.3e-06
1.1e-09
9.9e-06
4.8e-09
1.6e-07
1.1e-08
1.4e-05
1.3e-08
2.4e-08
3.1e-10
6.9e-09
2.4e-10
1.8e-08
1.9e-10
1.8e-01
1.8e-10
1.9e-07
1.2e-11
2.5e-06

**Supplementary Figure 2: APNet unveils the top 10 critical drivers in Severe COVID-19 patients across 3 Olink datasets. (A-C)** GSEA-like plots depicting the top 5 positive (red) and top 5 negative (blue) drivers and their related targets from Differential Activity analysis through NetBID2 workflow, throughout 3 Olink datasets [MGH (A), Mayo (B), and Stanford (C)] along with heatmaps showing functional clustering of corresponding drivers with their biological pathways.

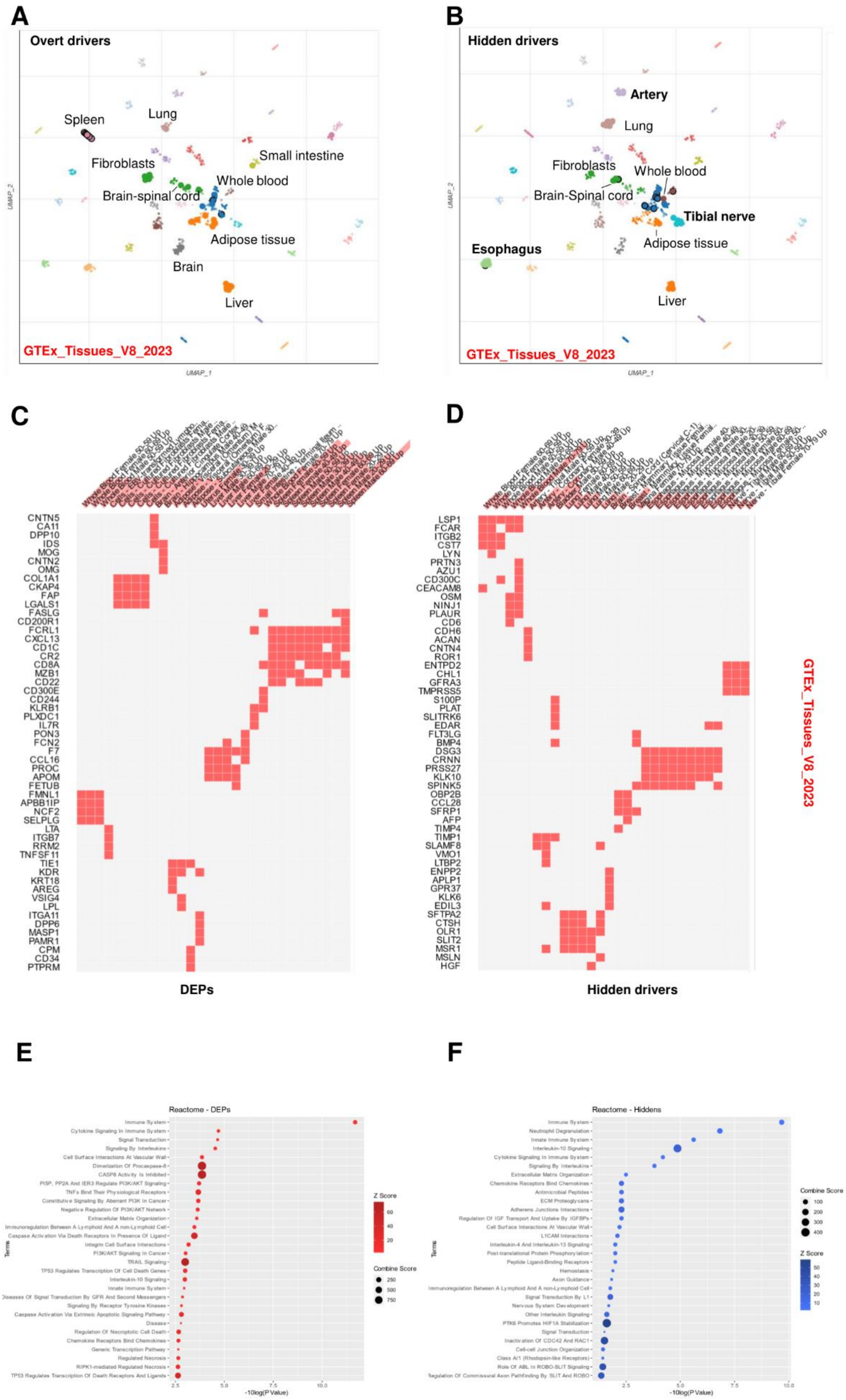
A
Overt drivers
Spleen
Lung
Fibroblasts
Small intestine
Whole blood
Brain-spinal cord
Adipose tissue
Brain
Liver
GTEx_Tissues_V8_2023
UMAP_1
UMAP_2
B
Hidden drivers
Artery
Lung
Fibroblasts
Whole blood
Brain-Spinal cord
Tibial nerve
Esophagus
Adipose tissue
Liver
GTEx_Tissues_V8_2023
UMAP_1
UMAP_2
C
CNTN5
CA11
DPP10
IDS
MOG
CNTN2
OMG
COL1A1
CKAP4
FAP
LGALS1
FASLG
CD200R1
FCRL1
CXCL13
CD1C
CR2
CD8A
MZB1
CD22
CD300E
CD244
KLRB1
PLXDC1
IL7R
PON3
FCN2
F7
CCL16
PROC
APOM
FETUB
FMNL1
APBB1IP
NCF2
SELPLG
LTA
ITGB7
RRM2
TNFSF11
TIE1
KDR
KRT18
AREG
VSIG4
LPL
ITGA11
DPP6
MASP1
PAMR1
CPM
CD34
PTPRM
DEPs
D
LSP1
FCAR
ITGB2
CST7
LYN
PRTN3
AZU1
CD300C
CEACAM8
OSM
NINJ1
PLAUR
CD6
CDH6
ACAN
CNTN4
ROR1
ENTPD2
CHL1
GFRA3
TMPRSS5
S100P
PLAT
SLITRK6
EDAR
FLT3LG
BMP4
DSG3
CRNN
PRSS27
KLK10
SPINK5
OBP2B
CCL28
SFRP1
AFP
TIMP4
TIMP1
SLAMF8
VMO1
LTBP2
ENPP2
APLP1
GPR37
KLK6
EDIL3
SFTPA2
CTSH
OLR1
SLIT2
MSR1
MSLN
HGF
Hidden drivers
GTEx_Tissues_V8_2023
E
Reactome - DEPs
Terms
-10log(P Value)
Z Score
Combine Score
F
Reactome - Hiddens
Terms
-10log(P Value)
Combine Score
Z Score

**Supplementary Figure 3**. **The APNet pipeline has revealed COVID-19 hidden drivers that are specific to different tissues and distinct signaling pathways. (A-B)** UMAP plots depicting cellular enrichment analysis for DEPs drivers (common differential expressed proteins, DEPs) (A) and hidden drivers (B) among the three Olink proteomic studies based on Enrichr GTEx_Tissues_V8_2023 database **(C-D)** Heatmaps depicting proteins and cell types from (A) and (B) respectively. **(E-F)** Bubble plots depicting over-representation analysis based on WikiPathways 2021 for DEPs and hidden drivers of COVID-19 severity.

A

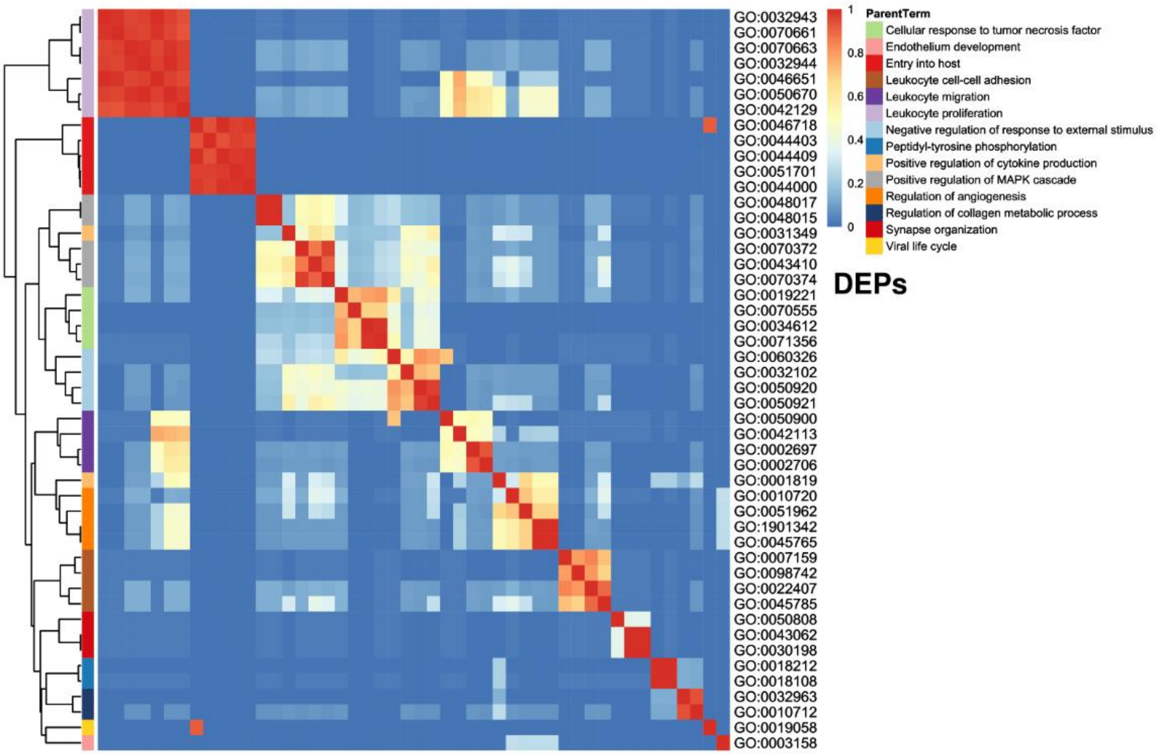
GO:0032943
GO:0070661
GO:0070663
GO:0032944
GO:0046651
GO:0050670
GO:0042129
GO:0046718
GO:0044403
GO:0044409
GO:0051701
GO:0044000
GO:0048017
GO:0048015
GO:0031349
GO:0070372
GO:0043410
GO:0070374
GO:0019221
GO:0070555
GO:0034612
GO:0071356
GO:0060326
GO:0032102
GO:0050920
GO:0050921
GO:0050900
GO:0042113
GO:0002697
GO:0002706
GO:0001819
GO:0010720
GO:0051962
GO:1901342
GO:0045765
GO:0007159
GO:0098742
GO:0022407
GO:0045785
GO:0050808
GO:0043062
GO:0030198
GO:0018212
GO:0018108
GO:0032963
GO:0010712
GO:0019058
GO:0003158
1
0.8
0.6
0.4
0.2
0
ParentTerm
Cellular response to tumor necrosis factor
Endothelium development
Entry into host
Leukocyte cell-cell adhesion
Leukocyte migration
Leukocyte proliferation
Negative regulation of response to external stimulus
Peptidyl-tyrosine phosphorylation
Positive regulation of cytokine production
Positive regulation of MAPK cascade
Regulation of angiogenesis
Regulation of collagen metabolic process
Synapse organization
Viral life cycle
DEPs

B

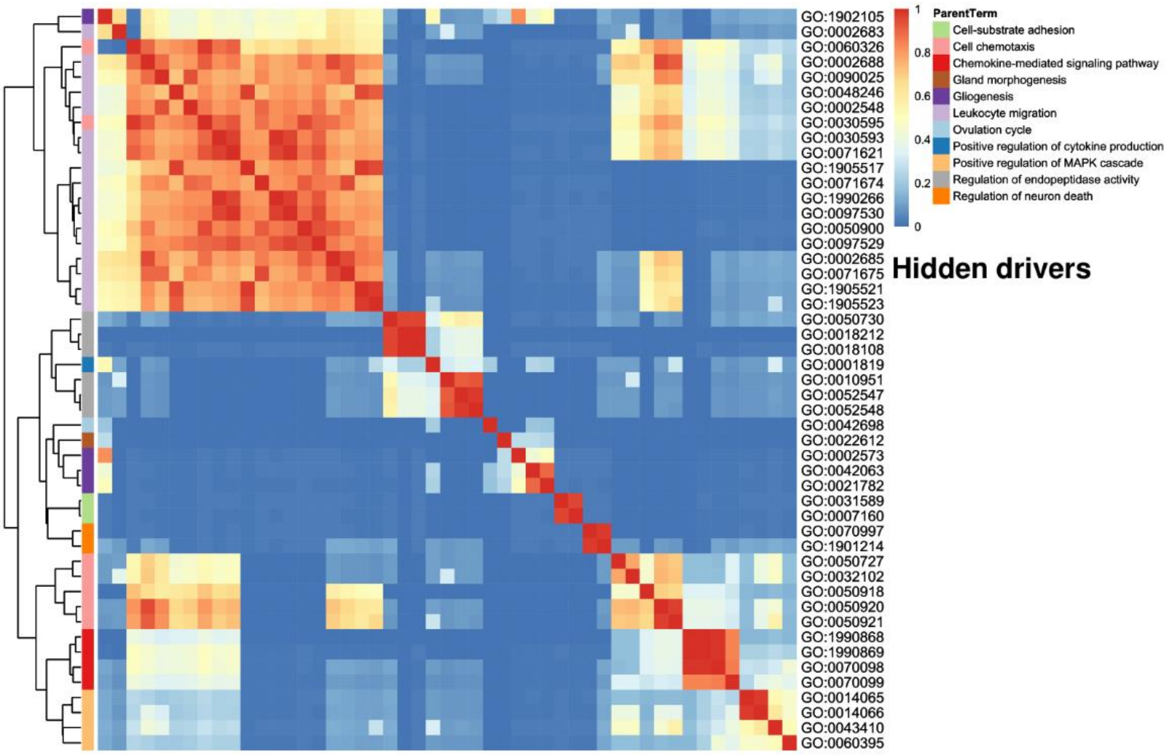
GO:1902105
GO:0002683
GO:0060326
GO:0002688
GO:0090025
GO:0048246
GO:0002548
GO:0030595
GO:0030593
GO:0071621
GO:1905517
GO:0071674
GO:1990266
GO:0097530
GO:0050900
GO:0097529
GO:0002685
GO:0071675
GO:1905521
GO:1905523
GO:0050730
GO:0018212
GO:0018108
GO:0001819
GO:0010951
GO:0052547
GO:0052548
GO:0042698
GO:0022612
GO:0002573
GO:0042063
GO:0021782
GO:0031589
GO:0007160
GO:0070997
GO:1901214
GO:0050727
GO:0032102
GO:0050918
GO:0050920
GO:0050921
GO:1990868
GO:1990869
GO:0070098
GO:0070099
GO:0014065
GO:0014066
GO:0043410
GO:0060395
1
0.8
0.6
0.4
0.2
0
ParentTerm
Cell-substrate adhesion
Cell chemotaxis
Chemokine-mediated signaling pathway
Gland morphogenesis
Gliogenesis
Leukocyte migration
Ovulation cycle
Positive regulation of cytokine production
Positive regulation of MAPK cascade
Regulation of endopeptidase activity
Regulation of neuron death
Hidden drivers

**Supplementary Figure 4. Hidden drivers of COVID-19 severity are more associated with perturbed lymphocyte trafficking and neurogenesis impairments than DEPs.** (A-B) Similarity heatmaps for GO:BP terms from the GeneKitr package after enrichment of joint DEPs (A) and hidden (B) drivers of COVID-19 severity, across the three Olink proteomic studies.

A

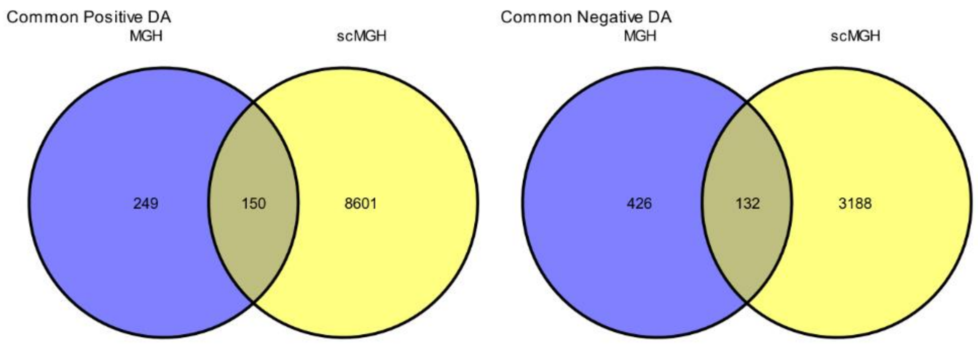
Common Positive DA
MGH
scMGH
249
150
8601
Common Negative DA
MGH
scMGH
426
132
3188

B

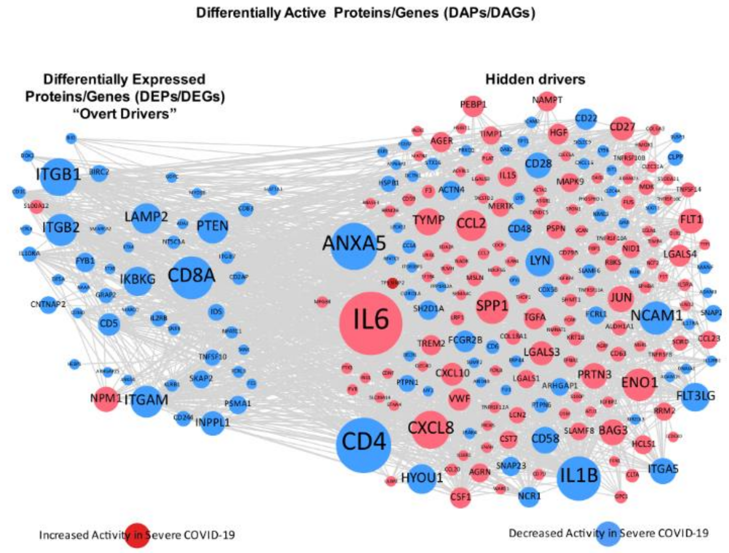
Differentially Active Proteins/Genes (DAPs/DAGs)
Differentially Expressed Proteins/Genes (DEPs/DEGs) "Overt Drivers"
Hidden drivers
ITGB1
ITGB2
LAMP2
PTEN
CD8A
IKBKG
ANXA5
IL6
CD4
IL1B
TYMP
CCL2
SPP1
CXCL8
NCAM1
ENO1
FLT3LG
HYOU1
ITGAM
NPM1
INPPL1
Increased Activity in Severe COVID-19
Decreased Activity in Severe COVID-19

C

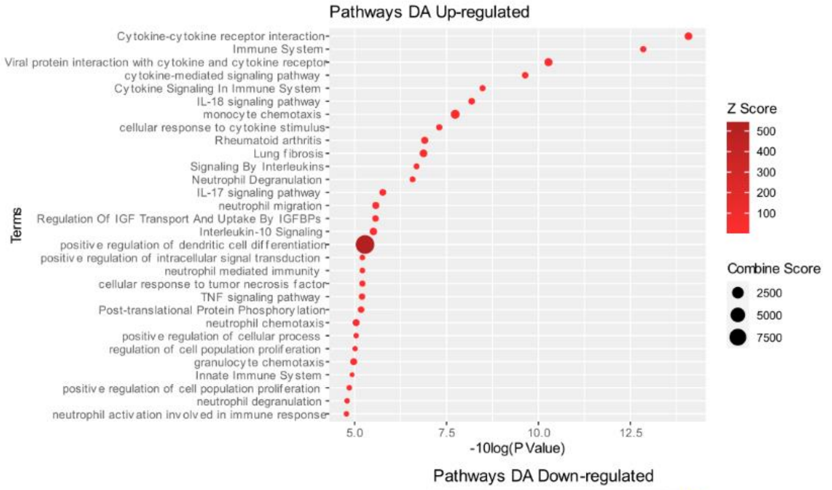
Pathways DA Up-regulated
Cytokine-cytokine receptor interaction
Immune System
Viral protein interaction with cytokine and cytokine receptor
cytokine-mediated signaling pathway
Cytokine Signaling In Immune System
IL-18 signaling pathway
monocyte chemotaxis
cellular response to cytokine stimulus
Rheumatoid arthritis
Lung fibrosis
Signaling By Interleukins
Neutrophil Degranulation
IL-17 signaling pathway
neutrophil migration
Regulation Of IGF Transport And Uptake By IGFBPs
Interleukin-10 Signaling
positive regulation of dendritic cell differentiation
positive regulation of intracellular signal transduction
neutrophil mediated immunity
cellular response to tumor necrosis factor
TNF signaling pathway
Post-translational Protein Phosphorylation
neutrophil chemotaxis
positive regulation of cellular process
regulation of cell population proliferation
granulocyte chemotaxis
Innate Immune System
positive regulation of cell population proliferation
neutrophil degranulation
neutrophil activation involved in immune response
Terms
5.0
7.5
10.0
12.5
-10log(P Value)
Z Score
500
400
300
200
100
Combine Score
2500
5000
7500

D

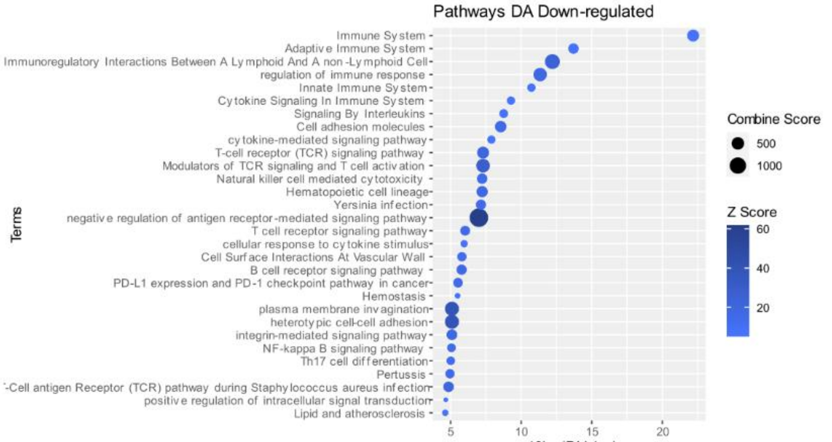
Pathways DA Down-regulated
Immune System
Adaptive Immune System
Immunoregulatory Interactions Between A Lymphoid And A non-Lymphoid Cell
regulation of immune response
Innate Immune System
Cytokine Signaling In Immune System
Signaling By Interleukins
Cell adhesion molecules
cytokine-mediated signaling pathway
T-cell receptor (TCR) signaling pathway
Modulators of TCR signaling and T cell activation
Natural killer cell mediated cytotoxicity
Hematopoietic cell lineage
Yersinia infection
negative regulation of antigen receptor-mediated signaling pathway
T cell receptor signaling pathway
cellular response to cytokine stimulus
Cell Surface Interactions At Vascular Wall
B cell receptor signaling pathway
PD-L1 expression and PD-1 checkpoint pathway in cancer
Hemostasis
plasma membrane invagination
heterotypic cell-cell adhesion
integrin-mediated signaling pathway
NF-kappa B signaling pathway
Th17 cell differentiation
Pertussis
positive regulation of intracellular signal transduction
Lipid and atherosclerosis
Terms
5
10
15
20
Combine Score
500
1000
Z Score
60
40
20

Supplementary Figure 5. **APNet reveals common drivers of severity across MGH peripheral blood and scRNA-seq data. (A)** Venn diagrams showing overlapping differentially active genes (DAGs) from MGH PBMC scRNA-seq data with differentially active proteins (DAPs) from the MGH plasma proteomics. **(B)** STRINGdb protein-protein interaction networks for joint DEPs and hidden drivers of severity across the two studies (STRINGdb score > 0.4). The size of the nodes is analogous to the centrality of each protein/driver (BetweenessCentrality algorithm) and the colour denotes perturbational direction (red for increased, blue for decreased). **(C-D)** Bubble plots depicting over-representation analysis based on the Enrichr Knowledge Graph (Wikipathways 2021, Reactome, GO:BP, KEGG) for joint drivers with increased (C) and decreased activity (D) in severe COVID-19 cases, among the three Olink plasma proteomic studies.

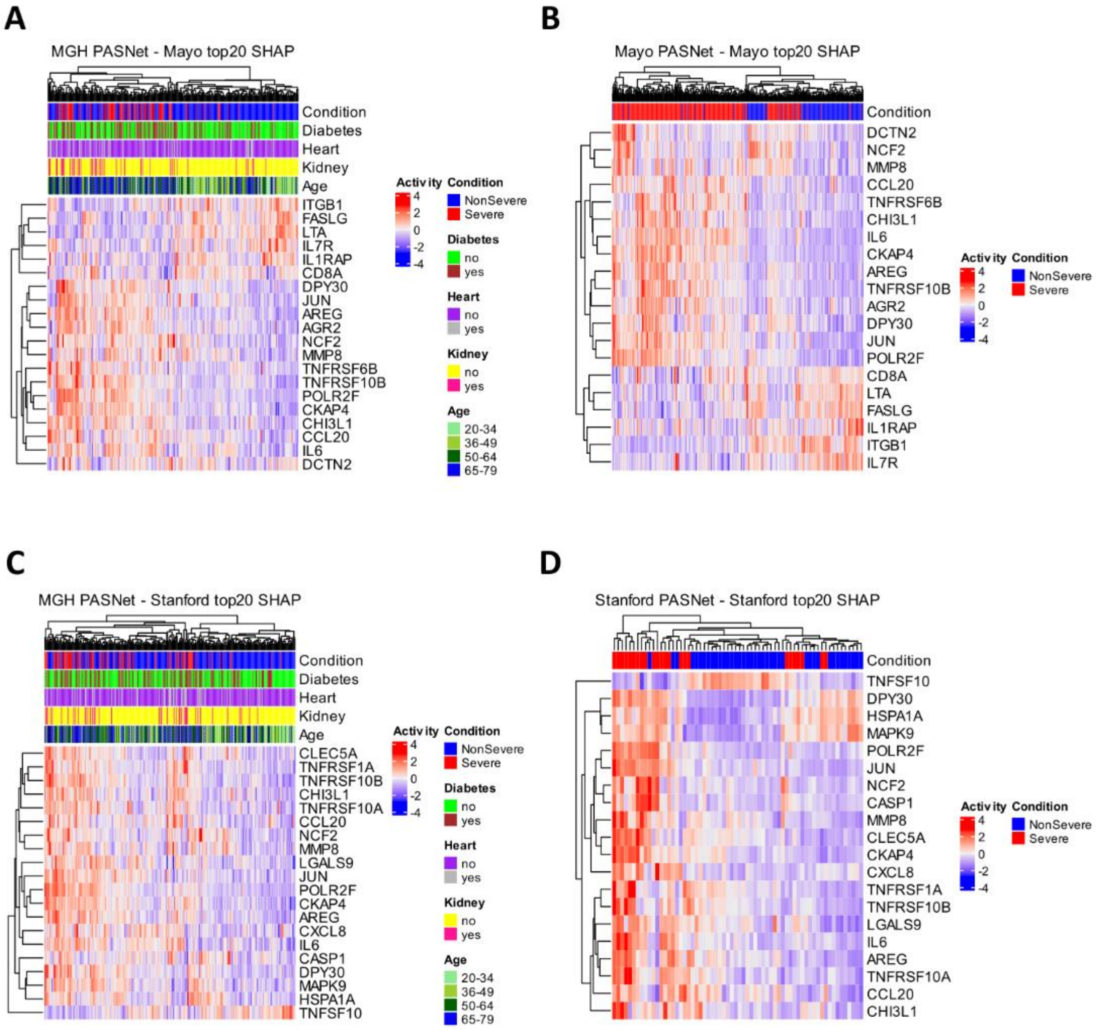
A
MGH PASNet - Mayo top20 SHAP
Condition
Diabetes
Heart
Kidney
Age
ITGB1
FASLG
LTA
IL7R
IL1RAP
CD8A
DPY30
JUN
AREG
AGR2
NCF2
MMP8
TNFRSF6B
TNFRSF10B
POLR2F
CKAP4
CHI3L1
CCL20
IL6
DCTN2
Activity
4
2
0
-2
-4
Condition
NonSevere
Severe
Diabetes
no
yes
Heart
no
yes
Kidney
no
yes
Age
20-34
36-49
50-64
65-79
B
Mayo PASNet - Mayo top20 SHAP
Condition
DCTN2
NCF2
MMP8
CCL20
TNFRSF6B
CHI3L1
IL6
CKAP4
AREG
TNFRSF10B
AGR2
DPY30
JUN
POLR2F
CD8A
LTA
FASLG
IL1RAP
ITGB1
IL7R
Activity
4
2
0
-2
-4
Condition
NonSevere
Severe
C
MGH PASNet - Stanford top20 SHAP
Condition
Diabetes
Heart
Kidney
Age
CLEC5A
TNFRSF1A
TNFRSF10B
CHI3L1
TNFRSF10A
CCL20
NCF2
MMP8
LGALS9
JUN
POLR2F
CKAP4
AREG
CXCL8
IL6
CASP1
DPY30
MAPK9
HSPA1A
TNFSF10
Activity
4
2
0
-2
-4
Condition
NonSevere
Severe
Diabetes
no
yes
Heart
no
yes
Kidney
no
yes
Age
20-34
36-49
50-64
65-79
D
Stanford PASNet - Stanford top20 SHAP
Condition
TNFSF10
DPY30
HSPA1A
MAPK9
POLR2F
JUN
NCF2
CASP1
MMP8
CLEC5A
CKAP4
CXCL8
TNFRSF1A
TNFRSF10B
LGALS9
IL6
AREG
TNFRSF10A
CCL20
CHI3L1
Activity
4
2
0
-2
-4
Condition
NonSevere
Severe

Supplementary Figure 6. **Benchmarking with PASNet outputs correlations between Severe and non-Severe COVID-19 patients' predictive drivers and clinical covariates.** Similar to Supplementary Figure 5, heatmaps were generated for the top 20 SHAP proteins in both the MGH - Mayo testing model, shown in (A) and (B), and the MGH - Stanford testing model, shown in (C) and (D), respectively.

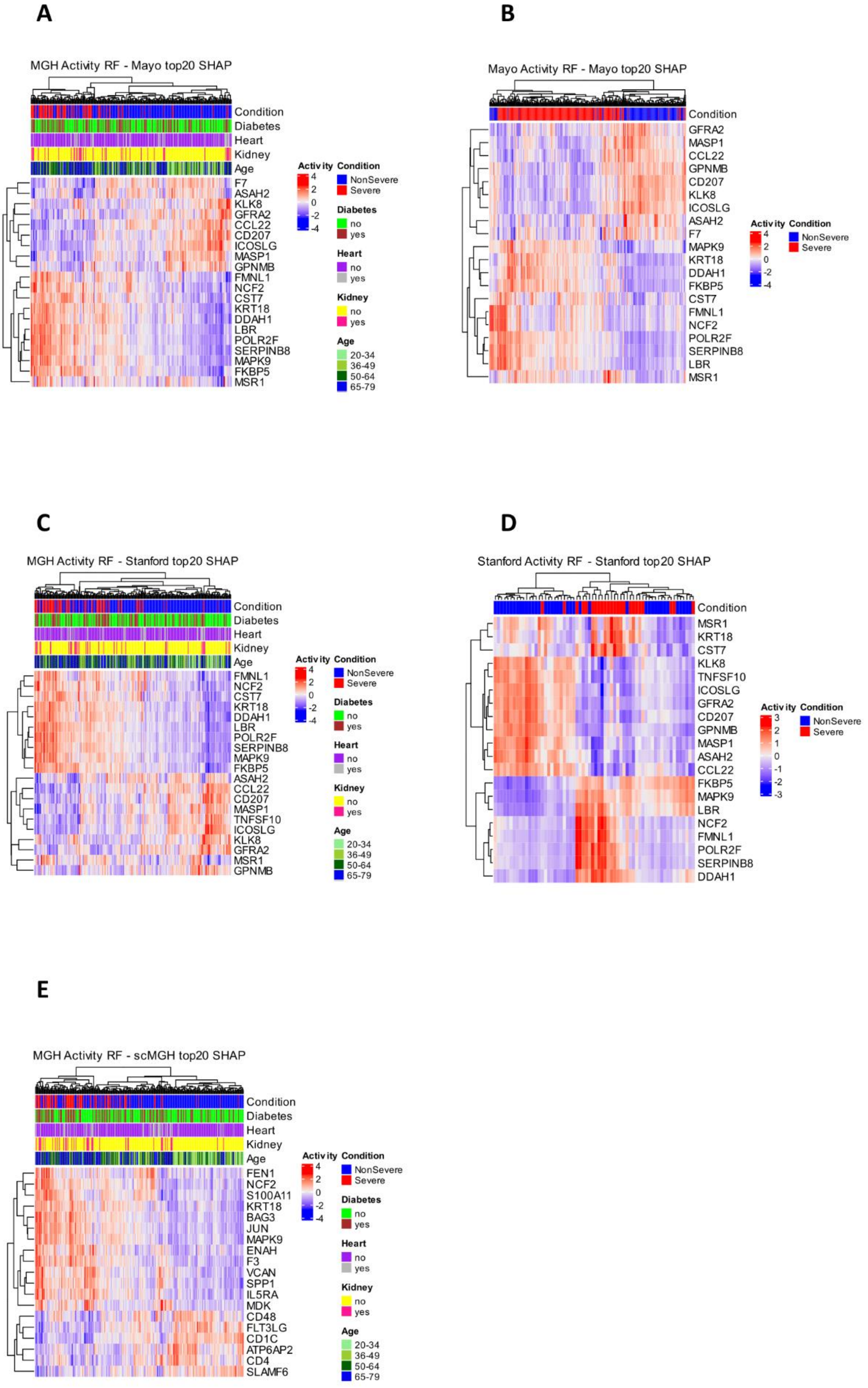
A
MGH Activity RF - Mayo top20 SHAP
Condition
Diabetes
Heart
Kidney
Age
F7
ASAH2
KLK8
GFRA2
CCL22
CD207
ICOSLG
MASP1
GPNMB
FMNL1
NCF2
CST7
KRT18
DDAH1
LBR
POLR2F
SERPINB8
MAPK9
FKBP5
MSR1
Activity
4
2
0
-2
-4
Condition
NonSevere
Severe
Diabetes
no
yes
Heart
no
yes
Kidney
no
yes
Age
20-34
36-49
50-64
65-79
B
Mayo Activity RF - Mayo top20 SHAP
Condition
GFRA2
MASP1
CCL22
GPNMB
CD207
KLK8
ICOSLG
ASAH2
F7
MAPK9
KRT18
DDAH1
FKBP5
CST7
FMNL1
NCF2
POLR2F
SERPINB8
LBR
MSR1
Activity
4
2
0
-2
-4
Condition
NonSevere
Severe
C
MGH Activity RF - Stanford top20 SHAP
Condition
Diabetes
Heart
Kidney
Age
FMNL1
NCF2
CST7
KRT18
DDAH1
LBR
POLR2F
SERPINB8
MAPK9
FKBP5
ASAH2
CCL22
CD207
MASP1
TNFSF10
ICOSLG
KLK8
GFRA2
MSR1
GPNMB
Activity
4
2
0
-2
-4
Condition
NonSevere
Severe
Diabetes
no
yes
Heart
no
yes
Kidney
no
yes
Age
20-34
36-49
50-64
65-79
D
Stanford Activity RF - Stanford top20 SHAP
Condition
MSR1
KRT18
CST7
KLK8
TNFSF10
ICOSLG
GFRA2
CD207
GPNMB
MASP1
ASAH2
CCL22
FKBP5
MAPK9
LBR
NCF2
FMNL1
POLR2F
SERPINB8
DDAH1
Activity
3
2
1
0
-1
-2
-3
Condition
NonSevere
Severe
E
MGH Activity RF - scMGH top20 SHAP
Condition
Diabetes
Heart
Kidney
Age
FEN1
NCF2
S100A11
KRT18
BAG3
JUN
MAPK9
ENAH
F3
VCAN
SPP1
IL5RA
MDK
CD48
FLT3LG
CD1C
ATP6AP2
CD4
SLAMF6
Activity
4
2
0
-2
-4
Condition
NonSevere
Severe
Diabetes
no
yes
Heart
no
yes
Kidney
no
yes
Age
20-34
36-49
50-64
65-79

**Supplementary Figure 7. Benchmarking with Random Forest outputs correlations between crucial predictive drivers and clinical covariates in Severe COVID-19.** Similar to Supplementary Figure 5, heatmaps were generated for the top 20 SHAP proteins in both the MGH - Mayo testing model, shown in (A) and (B), and the MGH - Stanford testing model, shown in (C) and (D), respectively. Also, The common 13 patients from MGH Olink and single cells MGH depicted into heatmap with the relevant covariates from MGH Olink datasets.

**A**

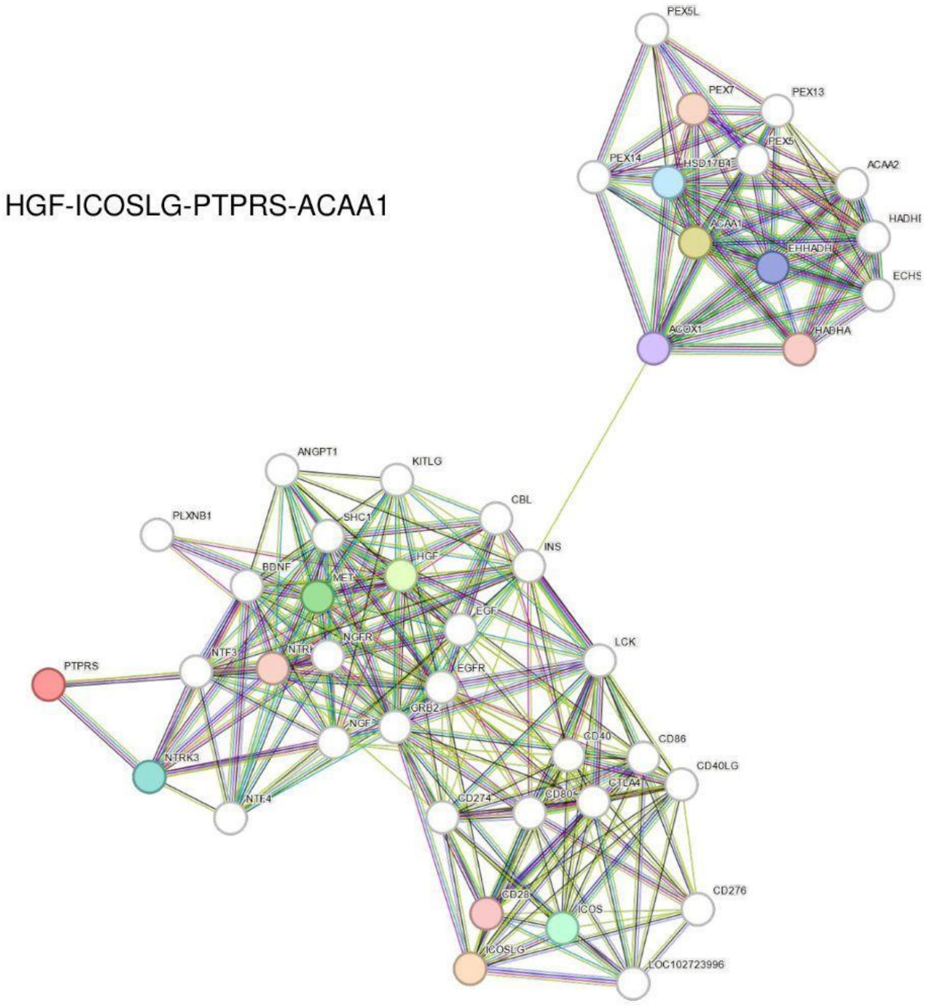
HGF-ICOSLG-PTPRS-ACAA1
PEX5L
PEX7
PEX13
PEX14
HSD17B4
PEX5
ACAA2
ACAA1
EHHADH
ECHS1
ACOX1
HADHA
ANGPT1
KITLG
PLXNB1
SHC1
CBL
INS
BDNF
MET
HGF
EGF
NGFR
PTPRS
NTF3
EGFR
LCK
NGF
GRB2
CD40
CD86
NTRK3
CD40LG
CTLA4
NTF4
CD274
CD276
CD28
ICOS
ICOSLG
LOC102723996

**B**

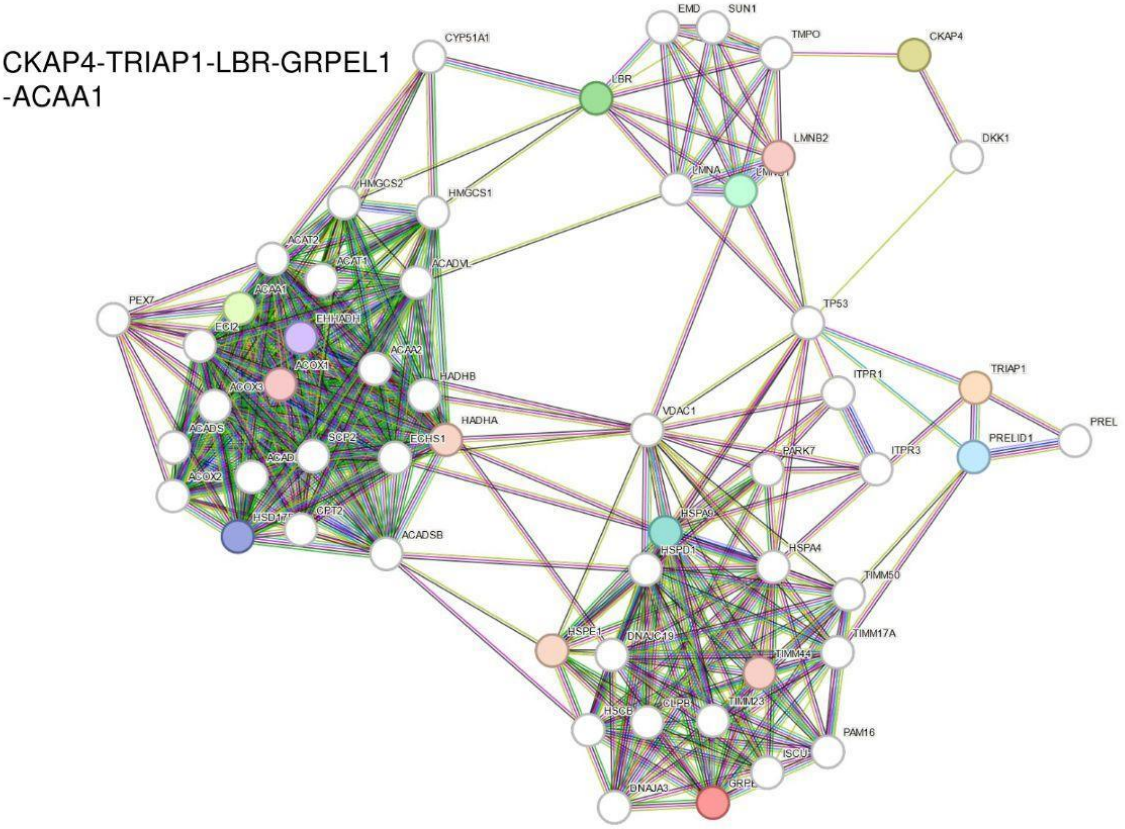
CKAP4-TRIAP1-LBR-GRPEL1
-ACAA1
EMD
SUN1
CYP51A1
TMPO
CKAP4
LBR
LMNB2
DKK1
LMNA
HMGCS2
HMGCS1
ACAT2
ACAT1
ACADVL
PEX7
ACAA1
EHHADH
TP53
ACAA2
ACOX1
ACOX3
HADHB
TRIAP1
ITPR1
ACADS
HADHA
VDAC1
PREL
SCP2
ECHS1
PARK7
ITPR3
PRELID1
ACOX2
CPT2
ACADSB
HSPA4
HSPD1
TIMM50
HSPE1
DNAJC19
TIMM17A
TIMM44
TIMM23
PAM16
ISCU
DNAJA3

**Supplementary Figure 8.** STRINGdb PPI networks of selected shortest paths from the MGH-weighted driver-pathway network as generated by APNet for HGF-ICOSLG-PTPRS-ACAA1 (A) and CKAP4-TRIAP1-LBR-GRPEL1-ACAA1 (B). Intermediate nodes were provided by STRINGdb (STRING score > 0.4).
